# Crest maturation at the cardiomyocyte surface contributes to a new late postnatal development stage that controls the diastolic function of the adult heart

**DOI:** 10.1101/2022.02.11.480042

**Authors:** Clément Karsenty, Céline Guilbeau-Frugier, Gaël Genet, Marie-Hélène Seguelas, Philippe Alzieu, Olivier Cazorla, Alexandra Montagner, Yuna Blum, Caroline Dubroca, Julie Maupoint, Blandine Tramunt, Marie Cauquil, Thierry Sulpice, Sylvain Richard, Silvia Arcucci, Remy Flores-Flores, Nicolas Pataluch, Romain Montoriol, Pierre Sicard, Antoine Deney, Thierry Couffinhal, Jean-Michel Sénard, Céline Galés

## Abstract

**RATIONALE:** In addition to its typical rod-shape, the mammalian adult cardiomyocyte (CM) harbors a unique lateral membrane surface architecture with periodic crests, relying on the presence of subsarcolemmal mitochondria (SSM) the role of which is still unknown.

**OBJECTIVE:** To investigate the development and functional role of CM crests during the postnatal period.

**METHODS AND RESULTS:** Electron/confocal microscopy and western-blot of left ventricular tissues from rat hearts indicated a late CM surface crest maturation, between postnatal day 20 (P20) and P60, as shown by substantial SSM swelling and increased claudin-5 cell surface expression. The P20-P60 postnatal stage also correlates with an ultimate maturation of the T-Tubules and the intercalated disk. At the cellular level, we identified an atypical CM hypertrophy characterized by an increase in long- and short-axes without myofibril addition and with sarcomere lateral stretching, indicative of lateral stretch-based CM hypertrophy. We confirmed the P20-P60 hypertrophy at the organ level by echocardiography but also demonstrated a transcriptomic program after P20 targeting all the cardiac cell populations. At the functional level, using Doppler echocardiography, we found that the P20-P60 period is specifically dedicated to the improvement of relaxation. Mechanistically, using CM-specific knock-out mice, we identified ephrin-B1 as a determinant of CM crest maturation after P20 controlling lateral CM stretch-hypertrophy and relaxation. Interestingly, while young adult *Efnb1*^CMspe−/−^ mice essentially show a relaxation impairment with exercise intolerance, they progressively switch toward heart failure with 100% KO mice dying after 13 months.

**CONCLUSIONS:** This study highlights a new late P20-P60 postnatal developmental stage of the heart in rodents during which the CM surface crests mature through an ephrin-B1-dependant mechanism and regulate the diastolic function. Moreover, we demonstrate for the first time that the CM crest architecture is cardioprotective.

## Introduction

The mammalian adult cardiomyocyte (CM) harbors a typical rod shape specifically dedicated to the function of the adult heart. Understanding the maturation of the adult rod-shaped CM can provide important insights for regenerative medicine, especially for the differentiation of hiPSC cells into fully mature rod-shaped adult CMs which still remains an obstacle. To date, the molecular events leading to the setting of the CM rod-shape during the post-natal stage are still unknown.

Indeed, despite fetal and adult CMs were extensively studied, the postnatal stage has only recently garnered an unprecedented interest only recently with the discovery of the potential of adult CMs to proliferate and regenerate the heart^1^. Thus, strategies in this field have focused on boosting this regenerative process by exploiting factors specifically involved in the early CM proliferation arrest that occurs during the postnatal period^2^. However, despite the postnatal maturation period characterizing the exit of the CM from the cell cycle, it also coincides with a morphogenesis step during which the CM switches from a proliferative rounded shape to a non-proliferative mature rod shape^3^. *In situ*, mammalian CMs stop dividing and start their morphogenesis maturation around postnatal day 7 (P7)^4,5^.

The postnatal morphogenesis stage of the CM relies on a polarization process that results in the asymmetric organization of the plasma membrane components underlying specific functions with an atypical basolateral polarity. It begins with a longitudinal elongation process of the CM to progressively evolve into the rod-shaped characteristic of the mature adult state. Contrary to epithelial cells harboring a basolateral and apical side, the adult CM lacks the apical side and looks like a barrel surrounded by a unique basal side with a basement membrane connecting the fibrillar extracellular matrix (ECM) and flanked by two intercalated disks (ID) involved in the CM-CM tight interactions, shaping the longitudinal alignment of myofibers within the tissue^6,7^. The ID phenocopies the lateral side of polarized epithelial cells with the presence of tight, adherent, gap junctions and desmosomes that play a specific role in the anchorage of the contractile myofilaments but also in the synchronization of the contraction. One architectural feature of the basal side of the adult CM, also called the lateral membrane, also relies on its important intracellular invaginations into transverse T-tubules, which play a key role in both action potential propagation and Ca^2+^ handling/excitation-contraction coupling^8^. Likewise, the extracellular side of the LM seems more complex than initially suspected, since, besides the presence of a costamere structure expressing classical transmembrane receptors (integrin and DGC dystroglycan-glycoprotein complex) that connect the ECM components to the intracellular myofibrils, we and others have reported the atypical presence of transmembrane proteins, claudin-5 and ephrin-B1^9,10^, more likely involved in cell-cell communication. This finding was unexpected since, contrary to the ID, the lateral membrane was so far viewed as a CM side lacking physical interactions with neighboring CMs. However, in recent years, using high-resolution nanoscale imaging, i.e. SCIM, AFM and MET, we and Gorelik’s lab have described a highly organized architecture of the lateral membrane on adult CMs with periodic crests^11–13^ filled with subsarcolemmal mitochondria (SSM) whose role is unknown. More recently, we provided evidence for the existence in the 3D cardiac tissue of intermittent lateral crest-crest contacts all along the lateral membrane through claudin5/claudin-5 tight junction interactions^12^, thus reconciling the presence of such proteins on the lateral face of the adult CM. Exactly when and how the lateral membrane crests of the CM maturate is completely unknown.

In this study, we investigated the maturation of the CM crests and their role during the postnatal period. We provide evidence for a late postnatal development stage, following the set-up of the rod shape, during which crests fully maturate through SSM swelling. We also show that this maturation step of the crests is ephrin-B1-dependent and specifically regulates the diastolic function of the adult heart.

## METHODS

The methods are described in detail in the Supplemental Material.

## RESULTS

### CM surface crests maturate late after postnatal day 20

We investigated crest maturation during the postnatal period on the left ventricle tissue of male rat hearts from different postnatal days (P0/birth, P5, P20, P60/young adult) using transmission electron microscopy (TEM) as previously described^12^. Surprisingly, at P0, while the contractile apparatus is disorganized in the neonatal CM, myofibrils are already orientated on the longitudinal axis of the cell with the first layer already anchored to the plasma membrane through Z-lines, and thus outlining a periodic crest-like architecture, more likely plasma membrane protrusions, which can yet be visualized all along the CM surface (**Figure 1A; Figure I in the Supplemental Material**). These immature and disordered crests already attempt to interact with crests from a neighboring CM. At this stage, no SSM could be observed. Then, crest maturation throughout the postnatal period occurs in two steps. A first early step, already completed by P5, relies on the maturation of the Z-lines, which delimits each sarcomere from the outer myofibril, allowing better visualization of the surface crest structure, concomitant to the myofilament alignment/organization that follows the morphological elongation of the CM (**Figure 1A, P5**). However, crests still display an unstructured morphology and their heights, directly correlated with the SSM number as previously described in adult CMs^12^, are small (“flat”-appearance) due to the lack or the presence of very small SSM, likely immature SSM (**Figure 1A**). By comparison, much larger but disorganized interfibrillar mitochondria (IFM) can be visualized in CMs at birth (P0) predominantly around CM nuclei (**Figure I in the Supplemental Material**), while they mature until P60 through both swelling (completed at P20) and alignment along the myofilaments (**Figure II in the Supplemental Material**). The crest immaturity persists until P20, a late stage of the postnatal period, while the CM has already implemented its rod shape (**Figure III in the Supplemental Material**) and completed its whole cytoarchitecture^14^, as indicated by the perfect alignment of sarcomere Z-lines between myofibril layers inferred from the α-actinin staining (**Figure IV in the Supplemental Material**). It is worth noting, that at P20, CMs display different morphologies with both rod-shaped and spindle-shaped CMs when compared with only rod-shaped CMs at P60 (**Figure III lower panels in the Supplemental Material**). Interestingly, a second but delayed maturation step of the surface crests occurs between P20 and the adult stage (P60) through substantial SSM swelling (**Figure 1A**), during which crest heights significantly expand up, correlating with an increase in the SSM number and area (**Figure 1B**). We further confirmed this maturation of the CM surface crests after P20 by analyzing the expression and localization of claudin-5, a tight-junctional protein that we previously described as a determinant of the lateral crest-crest interactions between the LM of neighbouring CMs^12^. While claudin-5 protein expression is progressively induced in the cardiac tissue at birth (P0), reaching its maximal expression at P10 (**Figure 1C, left panel**), complete localization of the protein at the lateral membrane of the CM occurs only between P20 and P60 (**Figure 1C, right panel**), most likely in agreement with the implementation of claudin-5/claudin-5-dependent interactions necessary to clip crests from neighboring CMs that we previously described at the adult stage ^12^. In line with this mechanism, atypical tight junctions connecting crests from the lateral face of neighbor CMs could be observed only in the P60 adult stage (**Figure V in the Supplemental Material**). Remarkably, the maturation of the CM lateral surface occurring between P20 and P60, at least in part through the setting of crest-crest interactions, occurs concomitantly with an ultimate maturation of both the ID and the T-tubules, relying on a spatial reorganization of their specific components. Thus, while connexin 43 (gap junctions), desmoplakin 1/2 (desmosomes) and N-cadherin (Adherens junctions) are located on both the lateral membrane and the ID at P20, they fully relocalize to the ID at P60 (**Figure VI in the Supplemental Material**). Likewise, RyR and caveolin-3, T-tubule markers (see Methods), are still misaligned at P20 while a perfect alignment along the sarcomere Z-lines on the short CM axis can be observed at P60 (**Figure VII A, B in the Supplemental Material**). In agreement, the TT power (TT regularity) tends to increase after P20 while the TT periodicity (frequency) is already established at P20 (**Figure VII C and Methods in the Supplemental Material**).

**Figure 1.**
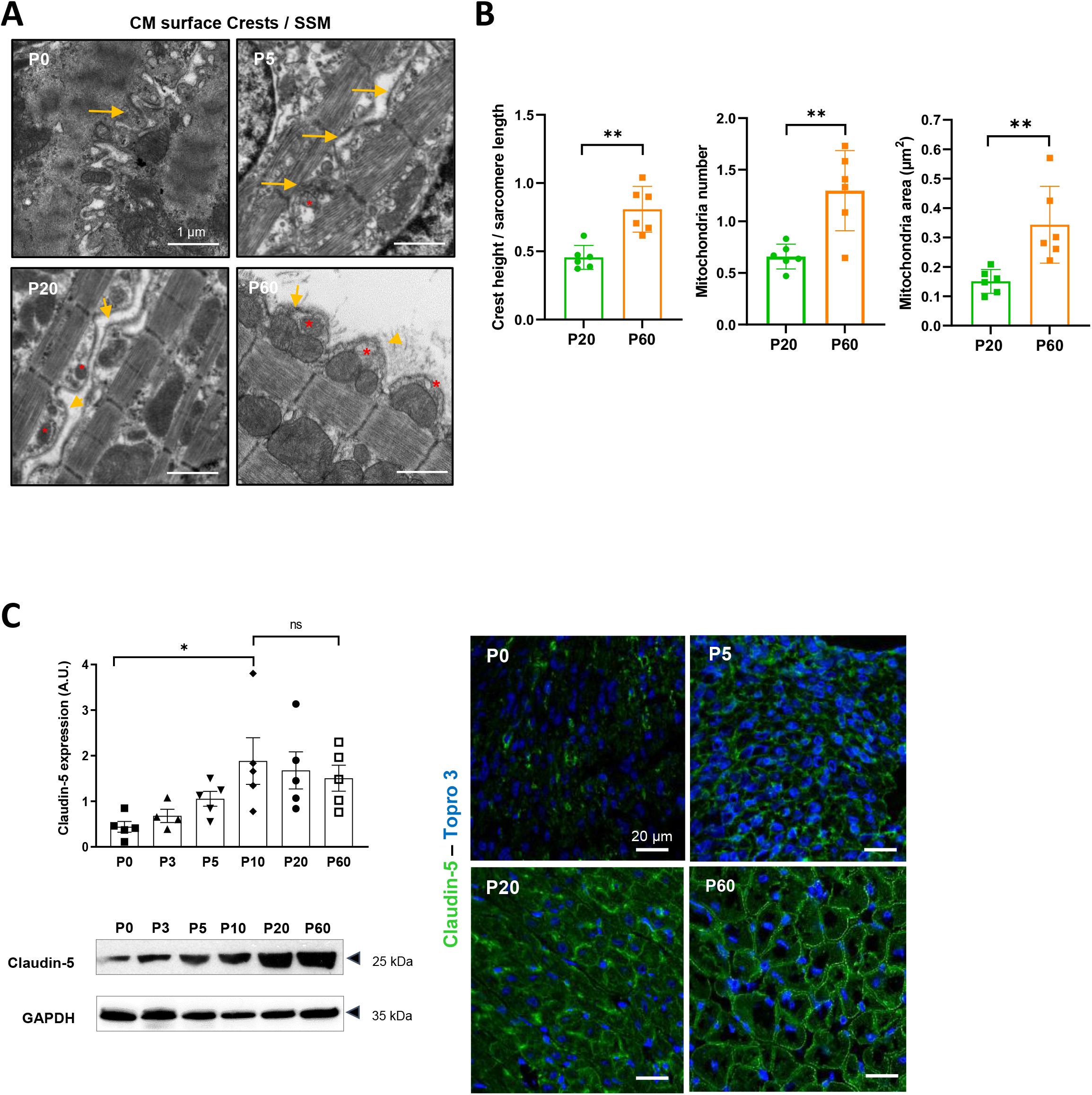
CM surface crests mature after postnatal day 20 (P20). **(A**) Transmission electron microscopy (TEM) micrographs showing representative CM surface crest relief (yellow arrows) and associated SSM (red stars) during rat postnatal maturation at postnatal day 0 (P0), 5 (P5), 20 (P20) and 60 (P60). (**B**) Quantification of crest heights / sarcomere length (*left panel*), SSM number / crest (*middle panel*) and SSM area (*right panel*) from TEM micrographs obtained from P20- or P60 rat hearts (P20 n=6, P60 n=6; 4 to 8 CMs/rat, ~ 70 crests/rat). (**C**) (*left panel*) Western-blot quantification of Claudin-5 protein expression in heart tissue from P0 to P60 old rats (upper panel) and representative immuno-blot (lower panel). (P0 n=5, P3 n=4; P5 n=5, P10 n=5, P20 n=5, P60 n=5); (*right panel*) Immunofluorescent localization of claudin-5 in heart cryo-sections from P0, P5, P20 and P60 rats. Data are presented as mean ± SD. Unpaired Student’s *t*-test for two group comparisons and one-way ANOVA for claudin-5 western-blot analysis, Tukey post-hoc test for six group comparisons with P0 as control. * *P* < 0.05, ** *P* < 0.01. ns, not significant.

Overall, these results indicate that the entire plasma membrane architecture of the CM maturates late, after P20, following the establishment of the rod shape.

### Evidence for a new late postnatal maturation stage of the CM and the mammalian heart dedicated to the development of the diastolic function

We next performed a more detailed analysis of the late P20/P60 maturation stage of the rodent heart.

At the cellular level, CMs from the left ventricle of rat hearts undergo significant hypertrophy between P20 and P60 as indicated by the substantial increase in their cross-sectional area (**Figure 2A**), which peaks at P45 (**Figure VIII in the Supplemental Material**), and in both their long and short axes (**Figure 2B and Figure IX in the Supplemental Material**). Similar late CM hypertrophy was observed in male mice (**Figure X in the Supplemental Material**). This P20-P60 CM hypertrophy was confirmed by echocardiography with a significant increase in the left ventricle posterior wall thickening (LVPWd) and cavity size (LVEDV, LVEDD) (**Figure 2C**), all indicative of an overall heart growth. Surprisingly, this physiological CM hypertrophy is atypical since it is not correlated with an expected increase in the myofilament compartment as indicated by the constant number of CM myofibrils between P20 and P60 (**Figure 2D**). This is corroborated by a marked decrease in the heart weight to body weight ratio during this period (**Figure 2E**), as previously reported^14^. Interestingly, we also noticed a significant and specific increase in the sarcomere heights with no variation of the sarcomere lengths (**Figure 2F**) which was inversely correlated with a decrease in the inter-lateral space between two CMs (**Figure 2G**), likely indicative of a lateral stretch of the CMs and a cardiac tissue compaction. Further supporting the CM lateral stretch that should distend the myofibrils, we observed larger distances between the thick myosin filaments at P60 than at P20 (**Figure XI in the Supplemental Material**). Taken together, these results highlight a new type of physiological cardiac hypertrophy that occurs during late postnatal development and that presumably relies, at least in part from the lateral vantage point, on the stretching of the CM lateral membrane.

**Figure 2.**
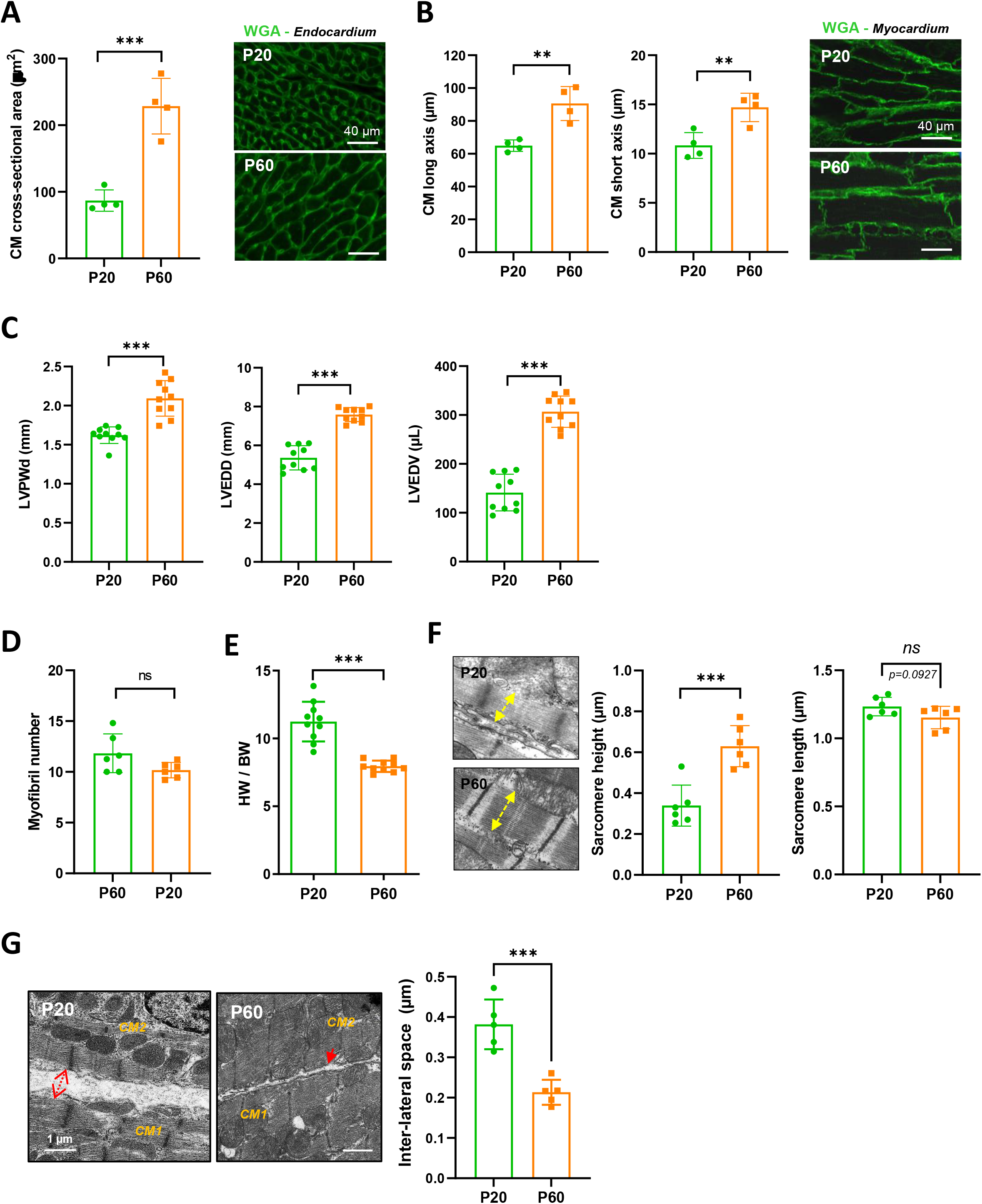
Evidence for a late postnatal maturation stage of the CM and the heart. (**A**) (*left panel*) CM area quantification from wheat-germ agglutinin (WGA)-stained heart cross-sections (endocardium) obtained from P20 and P60 rats (~ 400 CMs/rat; n=4 per group), as illustrated in the *right panel* (**B**) (*left panel*) CM long and short axis quantification from WGA-stained heart cross-sections (myocardium) obtained from P20 and P60 rats (~ 200 CMs/rat; n=4 per group) and illustrated in the *right panel*. (**C**) Analysis of echocardiography-based morphometry of hearts from P20 and P60 rats (P20 n=10, P60 n=10 rats). (**D**) Myofibril number quantified on the longitudinal CM axis from TEM micrographs of cardiac tissue from P20 or P60 rats (P20 n=6, P60 n=6; 4 to 8 CMs/rat). (**E**) Heart weight/body weight ratio of P20 or P60 rats (P20 n=10, P60 n=10). (**F)** (*left panel*) TEM micrographs showing representative sarcomere stretch (left arrows) from P20 to P60 rat; (*right panel*) Quantification of sarcomere height (P20 n=6, P60 n=6; 4 to 8 CMs/rat, ~ 35 sarcomeres/rat). (**G**) (*left panel*) TEM micrographs showing representative lateral membrane space between two neighboring CMs (red arrows) in cardiac tissue from P20 or P60 rats; (*right panel*) Quantification of the lateral membrane interspace (P20 n=6, P60 n=6; 4 to 8 CMs/rat, ~ 35 lateral interspaces/rat). Data are presented as mean ± SD. Unpaired student’s *t*-test for two group comparisons * *P* < 0.05, ** *P* < 0.01, *** *P* < 0.001. ns, not significant.

The existence of a new late postnatal maturation stage between P20 and P60 of the mammalian heart was confirmed by transcriptional analysis of left ventricular tissue from male mouse hearts. Volcano-plot analysis of the transcriptome and the heat map clearly show a significant difference between P20 and P60 heart gene expression (**Figure 3A, B**) with 1000 protein-coding genes that are up- or downregulated between P20 and P60 (p < 0.05, fold change > 1.5). The gene ontology (GO) enrichment analysis revealed significantly affected biological pathways (p < 0.05) (**Figure 3C and Figure XII in the Supplemental Material**) that are upregulated and mainly related to processes of the immune defence system, muscle cell differentiation, angiogenesis, positive regulation of cell death, different metabolisms including antioxidant defense, plasma membrane-related transport/signaling together with many pathways related to the nervous system development. By contrast, many ECM and developmental processes are downregulated. It is worth noting that some upregulated metabolic pathways relate to heart fuels other than fatty acids and glucose (tryptophan, glutamate, glycine/serine/threonine metabolisms), thus suggesting that the heart increases its metabolic flexibility between P20 and P60. We next performed gene clustering according to the cardiac cell populations established by Tucker et al*^15^* and assuming similar cardiac cell populations at P20 and P60 (**Methods in the Supplemental Material**). In line with the GO analysis, we found that the main downregulation of the transcriptome between P20 and P60 occurs primarily in the fibroblast populations known to be involved in the synthesis of the fibrillar ECM but also to a lesser extent in the CM populations (**Figure 3D**), consistent with the skeletal system and heart development pathways. By contrast, although all cardiac cells seem to be involved in the transcriptional maturation, the P20-P60 upregulated transcriptome largely referred to the ventricular CM populations when compared to the downregulated one (***Figure 3D***), in agreement with the evidence of a prominent muscle cell differentiation pathway depicted through the GO analysis. The existence of an important transcriptional maturation step of the CM during the P20-P60 postnatal window is further reinforced by the modulation of key CM-specific protein-encoding genes, especially several proteins from the contractile apparatus or regulating the contractile machinery at the lateral membrane (**Figure XIIIA in the Supplemental Material**). Another CM maturation also occurs at the metabolic level while the metabolic switch of the heart from glycolysis to fatty acid (FA) oxidation was already established during the early postnatal period^16^, with the remarkable upregulation of the *BDH1* gene that encodes the β-hydroxybutyrate dehydrogenase, the limiting mitochondrial enzyme for ketone body (KB) uptake during FA catabolism, but also to a lesser extent the upregulation of key actors in the glycolytic metabolism (*PFKFB2, SLC2A4*) (**Figure XIIIB in the Supplemental Material**).

**Figure 3.**
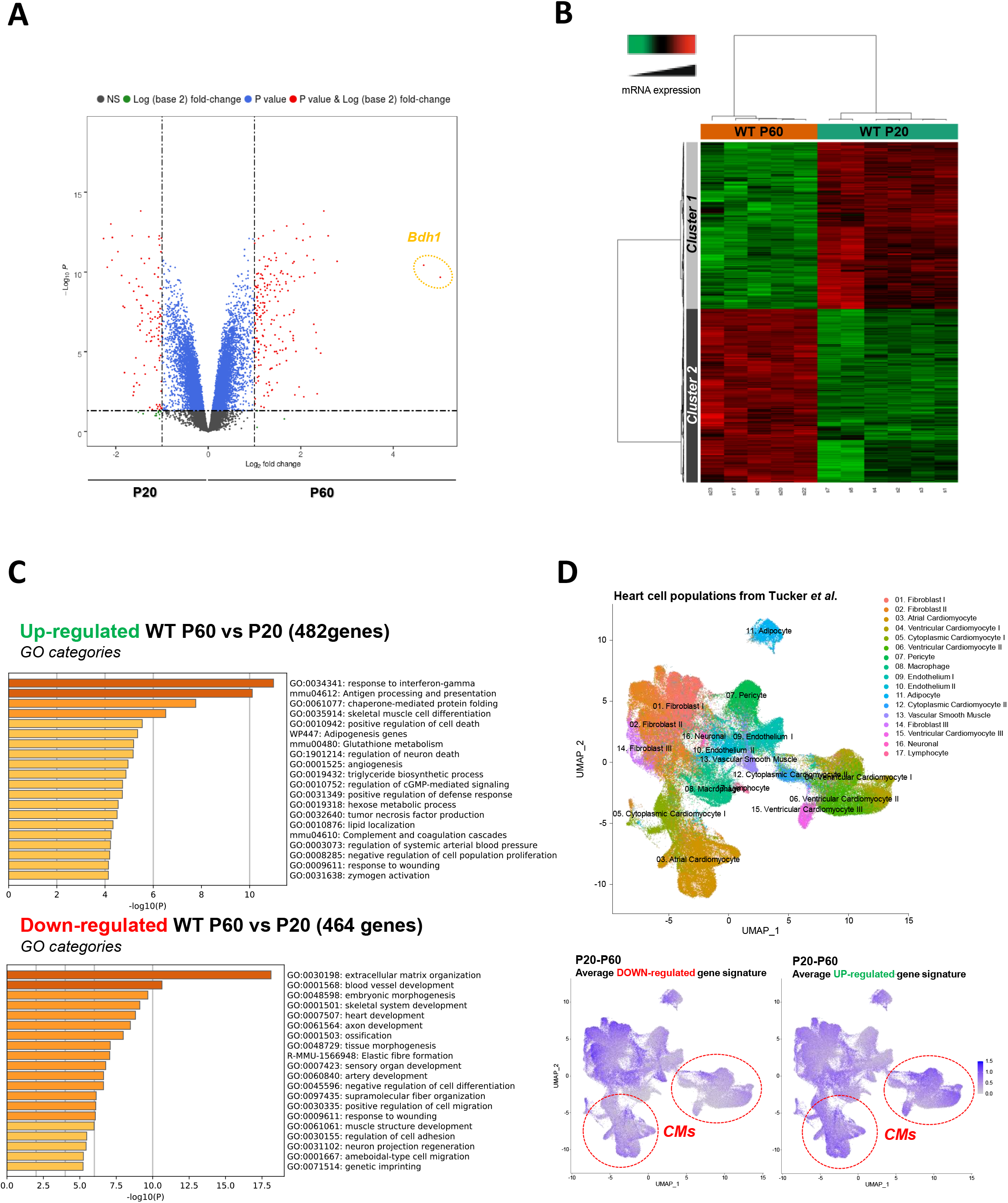
Late postnatal maturation stage between P20 and P60 of the mammalian heart confirmed by transcriptional analysis of left ventricular tissue. (**A**) Volcano plot of differences in gene expression between P20 and P60 mice. Colors indicate p < 0.05 and log (base 2) fold change > 2 (red), p < 0.05 and log (base 2) fold change < 2 (blue) and non-significant (NS) (black). (**B**) Heatmap presenting data from a microarray experiment performed with heart samples (P20 n=6, P60 n=5). Hierarchical clustering is also shown, which allows the definition of 2 gene clusters (p ≤ 0.05). (**C**) Gene ontology (GO) analysis of upregulated (*upper panel*) or downregulated (*lower panel*) genes between P60 and P20. The false discovery rate is provided for each category. (**D**) Uniform manifold approximation and projection (UMAP) plot displaying cellular diversity present in the human heart using Tucker *et al.’s* single-cell RNA-seq dataset. Each dot represents an individual cell. (*upper panel*) Colors correspond to the cell identity provided by the authors. The average expression of downregulated-(*left lower panel*) or up-regulated (*right lower panel*) gene signatures of left ventricles between P20 and P60 rats was calculated for each cell population and represented on the UMAP plot. Color key from gray to blue indicates relative expression level from low to high.

To better understand the functional impact of the P20-P60 maturation of the CM, we evaluated both the systolic and diastolic functions of rat hearts during this stage. Longitudinal echocardiographic evaluation reveals a similar left ventricular ejection fraction (LVEF) in both P20 and P60 rats (**Figure 4A**), indicating earlier maturation of the systolic function during the postnatal period. By contrast, we observed specific changes in the diastolic function between P20 and P60 as measured by non invasive Doppler imaging and showing a significant increase in passive filling (E/A) and an improvement in relaxation (*decrease in the isovolumic relaxation time (IVRT), increase in the e’/a’ and the early diastolic mitral annular tissue velocity e’, without change in the LV filling pressures E/e’*) (**Figure 4A**), most likely indicating that the diastolic function of the rat heart maturates during the late postnatal period, between P20 and P60. Further confirming the P20-P60 set-up of the adult diastolic function, while P20 and P60 rats display similar heart rates, invasive left ventricle catheter analysis shows increased systolic and diastolic blood pressure (SBP/DBP) as well as end-diastolic pressure (EDP) and an improvement in diastolic relaxation reflected by the dP/dtmin and the decreased time constant of isovolumic relaxation (Tau) (**Figure 4B**).

**Figure 4.**
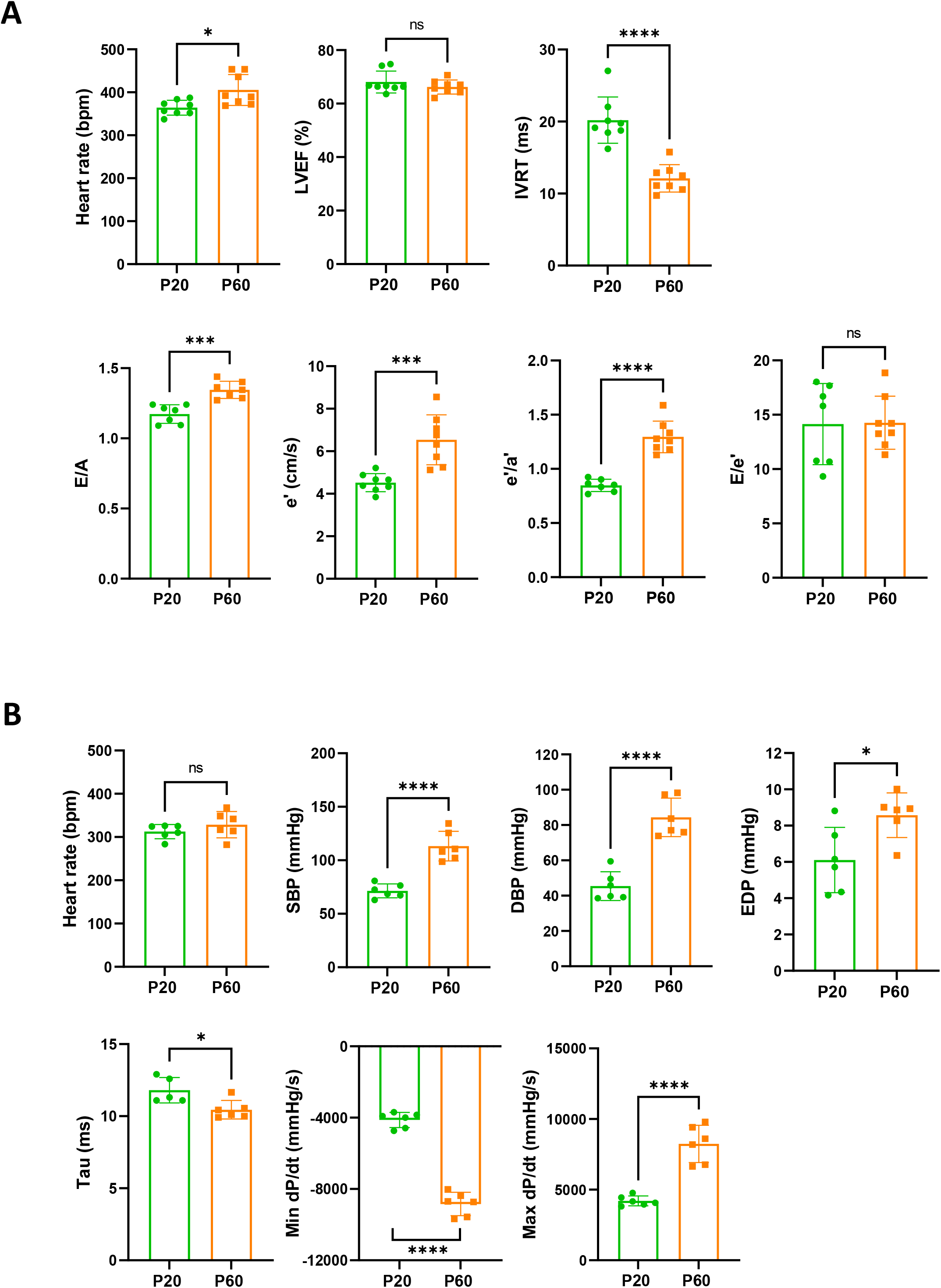
Late cardiac development is associated with the maturation of diastolic function. (**A**) Echocardiography performed on P20 and P60 rats using M-mode, B-mode, Doppler flow and tissular Doppler to measure systolic and diastolic function. Left ventricular ejection fraction (LVEF), isovolumetric relaxation time (IVRT), E/A ratio, e’ peak velocity, e’/a’ ratio and E/e’ ratio were measured (P20 n=8, P60 n=8). (**B**) Cardiac catheterization to assess the systolic and diastolic function of P20- or P60 rats. Systolic and diastolic blood pressure (SDB and DBP), end diastolic LV pressure (EDP), Tau and min and max dP/dt were measured (P20 n=6, P60 n=6). Data are presented as mean ± SD. Unpaired Student’s *t*-test for two group comparisons * *P* < 0.05, ** *P* < 0.01, *** *P* < 0.001, **** *P* < 0.0001 ns, not significant.

### *Efnb1*-specific knockdown in the CM impairs the late maturation of surface crests and the diastolic function

We have previously shown that the CM hypertrophy between P20 and P60 occurs concomitantly with the maturation of the crests that coat the whole CM lateral surface and, more specifically, with the implementation of the crest-crest interactions between neighbouring CMs. This observation raises questions about a specific contribution of the crest-crest interactions and the setting of intermittent tight junctions in the lateral stretch of the CM and the control of the diastole. Thus, we next sought to evaluate whether the maturation of the CM surface crests occurring after P20 could directly control the CM hypertrophy and the diastolic function of the heart.

For that purpose, we examined the role of ephrin-B1, a transmembrane protein that we previously identified as a new protein of the LM of the adult CM independent from the integrin or the dystroglycan complex, both connecting the ECM to the intracellular contractile machinery^9^. We showed that ephrin-B1 directly interacts with claudin-5 at the lateral membrane of the adult CM and controls its expression. Also, given the importance of claudin-5 in the setting of the crest-crest lateral interactions within the adult cardiac tissue^12^, we questioned about a potential role for ephrin-B1 as a crest determinant at the CM surface.

We first studied ephrin-B1 expression / localization in the cardiac tissue from rat hearts during the early postnatal development. As shown in **Figure 5A** and similarly to what we observed for claudin-5, ephrin-B1 reaches maximal expression very early during the postnatal period (P5). However, the complete trafficking of ephrin-B1 from the cytosol to the CM surface is only achieved at P60. To further explore the role of ephrin-B1, we took advantage of a knock-out (KO) mouse model harboring a CM-specific deletion of *Efnb1* that we previously described^9^. At the cellular level, the lack of ephrin-B1 in the CM partially impairs the P20-P60 physiological hypertrophy of the CM (**Figure 5B**), without modification of the myofibril number (**Figure 5C**). Consistent with ephrin-B1-specific expression at the lateral membrane, *Efnb1* deletion more specifically impedes the lateral stretch (short axis) of the CM but not the longitudinal one (**Figure 5D**) and accordingly leads to a decrease in both the sarcomere heights (**Figure XIVA in the Supplemental Material**) and the tissue compaction (increased lateral interspace) (**Figure XIVB in the Supplemental Material**). Of note, *Efnb1* deletion only partially prevents CM hypertrophy (partial CM short-axis elongation), most likely due to the other molecular events taking place at the LM during the P20-P60 stage, *i.e*. the interactions with the ECM (integrin, dystroglycan complex) that we previously showed to be independent of ephrin-B1^9^. At the lateral membrane level, P20 *Efnb1*^*CM−/−*^KO and WT mice harbor similar unstructured surface crests with immature SSM but only WT mice underwent crest maturation at P60 (**Figure 5E**), thus demonstrating a key role for ephrin-B1 in the postnatal maturation of the surface crest/SSM.

**Figure 5.**
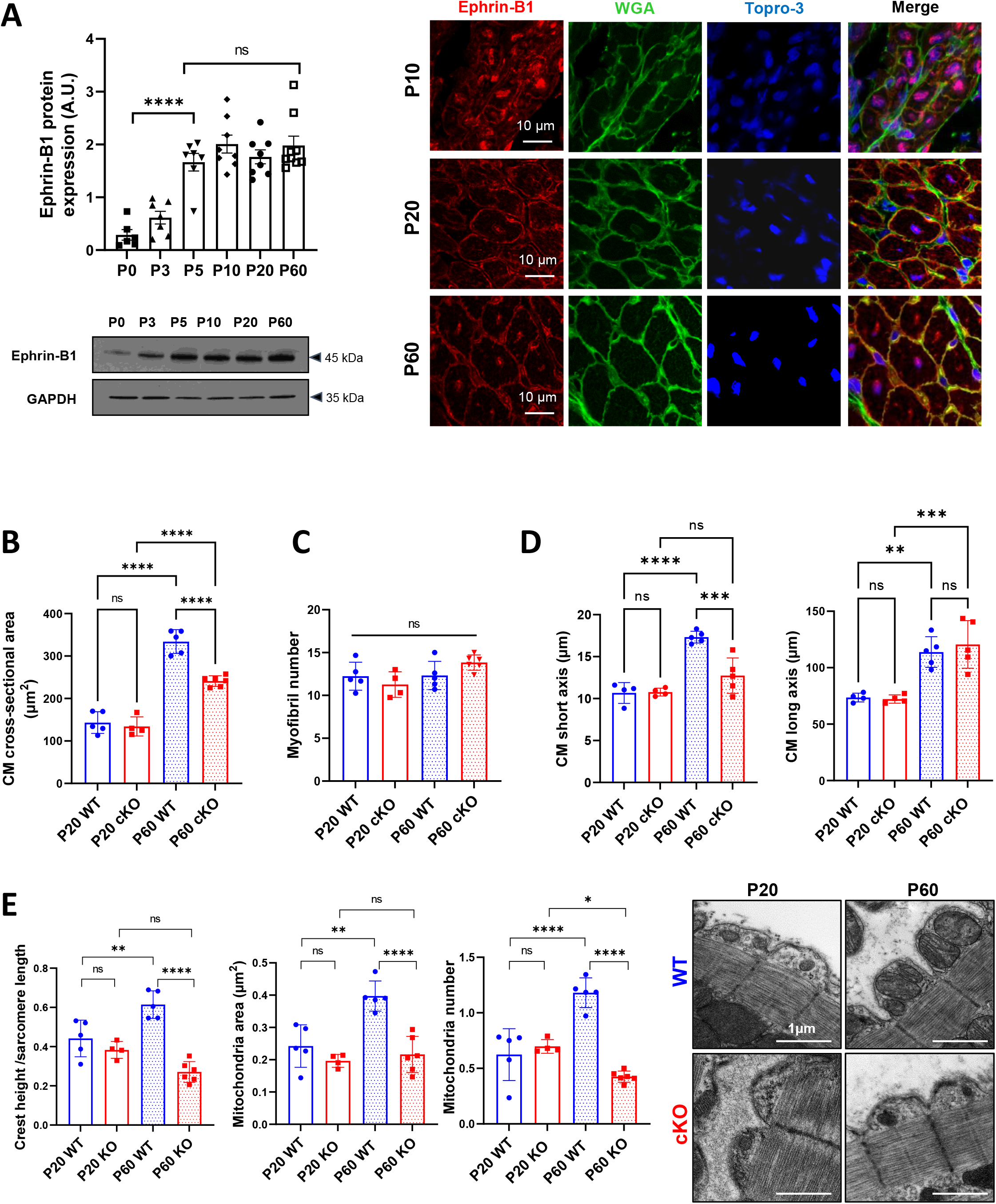
*Efnb1* specific knockdown in the CM impairs the late maturation of CM surface crests. (**A**) *(left panel*) Western-blot quantification of ephrin-B1 protein expression in heart tissue from P0 to P60 rats (upper) and representative immunoblot (lower) (P0 n=6, P3 n=7; P5 n=7, P10 n=8, P20 n=8, P60 n=8); (*right panel*) Immunofluorescent localization of ephrin-B1 (white arrows) in heart cryosection from P10, P20 and P60 old rats. At P10, ephrin-B1 (red) was mainly expressed in CM nuclei. At P20, ephrin-B1 is expressed at the CM lateral membrane but still highly in the CM cytoplasm while at P60, the protein is mainly expressed in the lateral membrane Nuclei were stained using topro-3 (blue). (**B**) CM area quantification from WGA-stained heart cross-sections (endocardium) obtained from P20 or P60 *Efnb1*^*CM-spe*^ KO and WT mice (~ 120 CMs/mice; WT P20 n = 5, cKO P20 n=4, WT P60 n=5, cKO n=6). (**C**) Myofibril number quantification from TEM micrographs of cardiac tissue from P20 or P60-old *Efnb1*^*CM-spe*^KO and WT mice (WT P20 n=5, cKO P20 n=4, WT P60 n=5, cKO n=6; 4 to 8 CMs/mouse). (**D**) CM long and short axis quantified from WGA-stained heart cross-sections (myocardium) obtained from P20 or P60 *Efnb1*^*CM-spe*^KO and WT mice (WT P20 n= 4, cKO P20 n=4, WT P60 n=5, cKO n=5; ~ 30 CMs/mouse). (**E**) Quantification of crest heights / sarcomere length (*left panel*), SSM area (*middle panel*) and SSM number / crest (*right panel*) from TEM micrographs obtained from P20- or P60 *Efnb1*^*CM-spe*^KO and WT mice (WT P20 n=5, cKO P20 n=4, WT P60 n=5, cKO n=5, ~ 60 crests per group) and illustrated in the *right panel*. One-way ANOVA, Tukey post-hoc test for 6 group comparisons with P0 as control. ****, P < 0.0001 for ephrin-B1 Western-blot analysis. Data are presented as mean ± SD. Unpaired Student’s *t*-test for two group comparisons or two-way ANOVA with Tukey post-hoc test for four group comparisons * *P* < 0.05, ** *P* < 0.01, *** *P* < 0.001, **** *P* <0.0001. ns, not significant.

We next assessed the role of ephrin-B1 in the cardiac function of young adult male mice (P60/2-months). We did not notice differences in heart rate and LVEF between 2-month-old WT and *Efnb1*^*CM-spe*^ KO mice (**Figure 6A**). However, and consistent with the role of ephrin-B1 in the lateral stretch-based hypertrophy of the CM, *Efnb1*^*CM−/−*^ KO mice display a significant decrease in LVPWd conversely to an increase in left ventricular internal diameter (Left ventricular internal diameter end diastole, LVIDd) and volume (LV end-diastolic volume, LVEDV) (**Figure 6A**), in agreement with our previous study^9^. Moreover, compared with WT mice, *Efnb1*^*CM−/−*^ KO mice display significantly elongated IVRT, increased left atrial volume (left atrial to aortic root ratio, LA/Ao) and decreased E/A as well as heterogeneous E/E’(**Figure 6A**), all indicative of impaired LV relaxation. Despite preservation of the LVEF, we also measured a significant decrease in ventricular global longitudinal strain (LV-GLS) in KO mice (**Figure 6A**) reflecting abnormal longitudinal systolic function. Diastolic dysfunction in KO mice was further confirmed by cardiac catheterization since 2-month-old *Efnb1*^*CM−/−*^ KO mice exhibit increased EDP and heterogeneous dP/dt but with similar relaxation time constants (tau) compared with age-matched WT mice (**Figure 6B**). The diastolic defects of *Efnb1*^*CM−/−*^ KO mice rely on a specific impairment of the CM relaxation since both the diastolic basal sarcomere length (SL) shortening and relaxation velocities are significantly reduced in isolated intact CMs from *Efnb1*^*CM−/−*^ KO compared to WT mice (**Figure 6C**). Furthermore, consistent with heart failure with preserved ejection fraction (HFpEF) phenotype in which CM contraction defects co-exist with the relaxation impairment ^17–19^, the auxotonic contraction indexed by the SL shortening as well as the SL shortening velocities are also significantly decreased in CMs from *Efnb1*^*CM−/−*^ KO compared to WT CMs (**Figure 6C**). Finally, *Efnb1*^*CM−/−*^ KO mice also display significant exercise intolerance compared with WT mice, as indicated by the significant decrease in maximum oxygen uptake (**Figure 6D**). Together, these results indicate that young adult male *Efnb1*^*CM−/−*^ KO mice recapitulate some features of clinical HFpEF, thus demonstrating a pivotal role for ephrin-B1 in the control of the diastolic function. In line with this assumption, a recent RNAseq study performed on ventricles from HFpEF, HFrEF (heart failure with reduced ejection fraction) and control patients reported a specific and more significant downregulation of the *Efnb1* gene in HFpEF than HFrEF patients (*p* = *9.10*^−14^ *HFpEF vs control, p* = *2.10*^*−5*^ *HFrEF versus control, p = 0.005 HFpEF versus HFrEF*)^20^.

**Figure 6.**
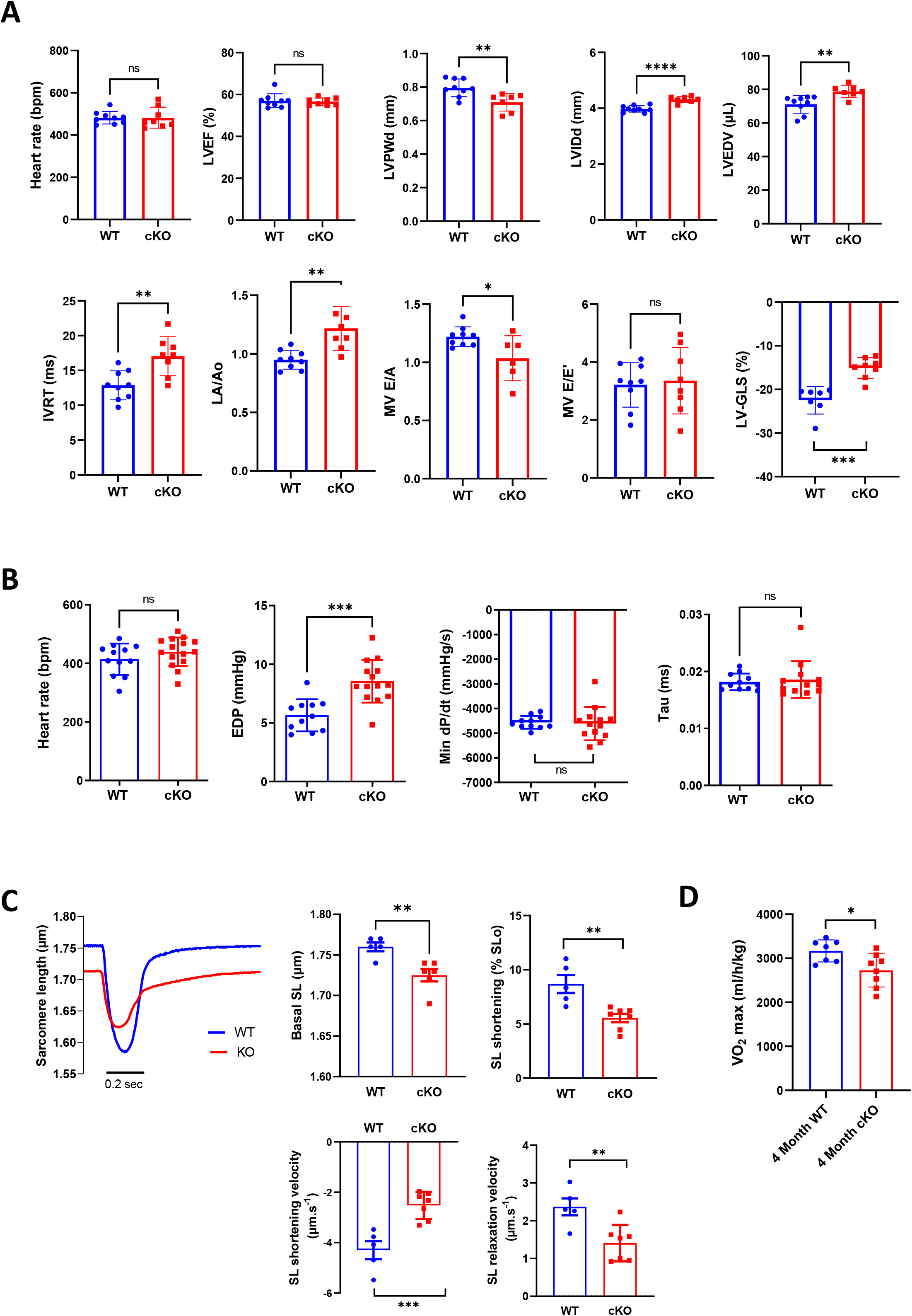
*Efnb1*-specific knockdown in the CM impairs diastolic function. (**A**) Echocardiography performed on P60 *Efnb1*^*CM−/−*^ KO or WT mice with 2D, TM, Doppler flow and tissular Doppler analysis to measure morphometry, systolic and diastolic function. Left ventricular ejection fraction (LVEF), interventricular septum wall thickness in diastole (IVSd), left ventricular internal diameter end diastole (LVIDd), left ventricular end-diastolic volume (LVEDV), isovolumetric relaxation time (IVRT), left atrium/aorta ratio (LA/Ao), E/A ratio, e’ peak velocity, e’/a’ ratio and E/e’ ratio were measured (WT n=9, cKO n=8). (**B**) Cardiac catheterization to assess diastolic function of P60 *Efnb1*^*CM−/−*^KO or WT mice, measuring end diastolic LV pressure (EDP), Tau, and min dP/dt (WT n=12, cKO n=14). (**C**) (*left panel*) Representative contraction evoked by electrical field stimulation as measured from sarcomere length (SL) shortening in isolated CMs (left ventricles) from P60 *Efnb1*^*CM-spe*^KO or WT mice; (*middle panel*) Basal sarcomere length (SL); (*right panel*) sarcomere length (SL) shortening during contraction; (*lower left panel*) sarcomere length shortening velocity; (*lower right panel*) Sarcomere length relaxation velocity (WT n=5, cKO n=7). (**D**) Treadmill exercise tolerance assay assessed by the VO2max measured from 4-month-old *Efnb1*^*CM-spe*^KO or WT mice (WT n=7, cKO n=8). Data are presented as mean ± SD. Unpaired Student’s *t*-test for two group comparisons * *P* < 0.05, ** *P* < 0.01, *** *P* < 0.001. **** *P* < 0.0001. ns, not significant.

More globally, these results demonstrate that ephrin-B1-dependent maturation of the CM surface crests during the P20-P60 postnatal period allows maturing the adult diastolic function.

### *Efnb1*^*CM−/−*^ mice switched progressively from HFpeF to HFrEF and all died at 14 months of age due to T-tubule disorganization

Finally, we monitored changes in the cardiac function of *Efnb1*^*CM−/−*^ KO and WT mice in medium and long term. Interestingly, while in young adulthood (2 months) *Efnb1*^*CM−/−*^ KO mice display LV diastolic dysfunction with preserved ejection fraction compared to WT mice, LVEF decreased progressively over time in these mice, which start developing moderate HFrEF at 9 months, evolving toward severe HFrEF by 13 months (**Figure 7A**) and 100% mortality after 15 months (**Figure 7B**). Accordingly, LVIDd and LVEDV significantly increase over time only in KO mice (**Figure 7A**) concurrently with the development of compensatory cardiac hypertrophy (**IVSd, Figure 7A, C, D**) and fibrosis around 12 months of age (**Figure 7E**). Corroborating the HF progression in *Efnb1*^*CM−/−*^ KO mice, while T-tubule (HFrEF marker) misalignment from Z-lines is already detectable but limited to discrete local regions in the cardiac tissue of some KO CMs at 2 months of age, this disorganization progressively spreads over time to all CMs after 12 months compared with WT mice (**Figure 7F**).

**Figure 7.**
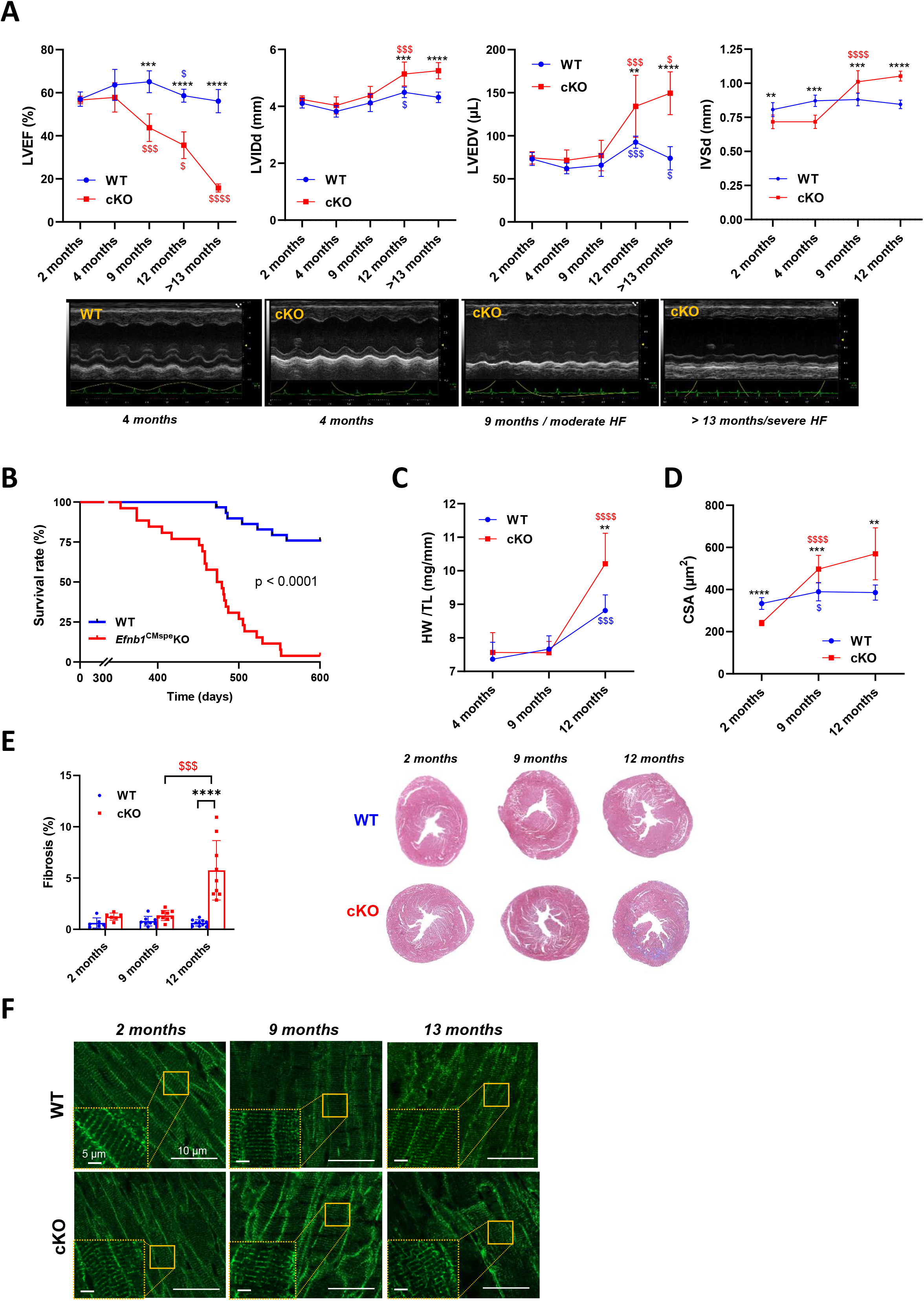
*Efnb1*^CM−/−^ mice switch progressively from HFpEF to HFrEF. (**A**) (*upper panels*) Longitudinal echocardiography to assess morphometry and systolic function of *Efnb1*^*CM−/−*^KO and WT mice over time. Left ventricular ejection fraction (LVEF), left ventricular internal diameter end diastole (LVIDd), left ventricular end diastolic volume (LVEDV), interventricular septum wall thickness in diastole (IVSd) were measured in 2-, 4-, 9-, 12- and > 13-month-old mice (WT 2 months n=9, 4 months n=4, 9 months n=9, 12 months n=9, >13 months n=9; cKO 2 months n=8, 4 months n=4, 9 months n=8, 12 months n=8, >13 months n=5). (*Lower panels*) Representative images of left ventricular M-mode echocardiography from WT compared to *Efnb1*^*CM−/−*^KO mice which show progressive LV dilatation and systolic function decline in the KO mice. (**B**) Kaplan-Meier survival plots for WT (blue) and *Efnb1*^*CM−/−*^KO mice (red). (Starting populations; WT n=26, cKO n=30) Survival analysis was performed by log-rank test. (**C**) Heart weight/tibia length ratios from WT and *Efnb1*^*CM−/−*^KO mice (WT 4 months n=6, 9 months n=8, 12 months n=8; cKO 4 months n=6, 9 months n=9, 12 months n=8). (**D**) CM area quantification from WGA-stained heart cross-sections (endocardium) from *Efnb1*^*CM−/−*^KO or WT mice (~ 120 CMs/mouse; WT 2 months n=5, 9 months n=9, 12 months n=8; cKO 2 months n=6, 9 months n=9, 12 months n=8). (**E**) (*left panel*) Cardiac fibrosis quantification from Masson’s trichrome staining of transverse sections from WT and *Efnb1*^*CM−/−*^KO mice hearts (2 months n=6, 9 months n=8, 12 months n=9) and (right panels) representative images. (**F**) Representative immunofluorescent staining of T-tubules (caveolin-3) in paraffin-embedded heart sections from *Efnb1*^*CM−/−*^KO and WT mice. Data are presented as mean ± SD. One-way ANOVA with Tukey post-hoc test for longitudinal group comparisons of WT or cKO mice ^$^ *P*<0.05, ^$$^ *P*<0.01, ^$$$^ *P*<0.001, ^$$$$^ *P*<0.0001. Unpaired Student’s *t*-test to compare WT and cKO groups. * *P* < 0.05, ** *P* < 0.01, *** *P* < 0.001, **** *P* < 0.0001. Only significant results are presented.

Collectively, these data indicate that the lack of ephrin-B1 in CM primarily impairs the diastolic function of young adult mice, which progressively switch toward a systolic defect with ageing.

## DISCUSSION

In this study, we describe a new developmental stage occurring in the late postnatal period between P20 and P60, during which LM crests maturate through SSM swelling and crest-crest interactions within the tissue, thus allowing CM lateral stretching and a global compaction of the cardiac tissue (**Figure 8**). We demonstrate that this mechanism is ephrin-B1-dependent and regulates the adult diastolic function. Taken together, our findings identify crest subdomains of the adult CM lateral surface as novel specific determinants of cardiac diastole.

**Figure 8.**
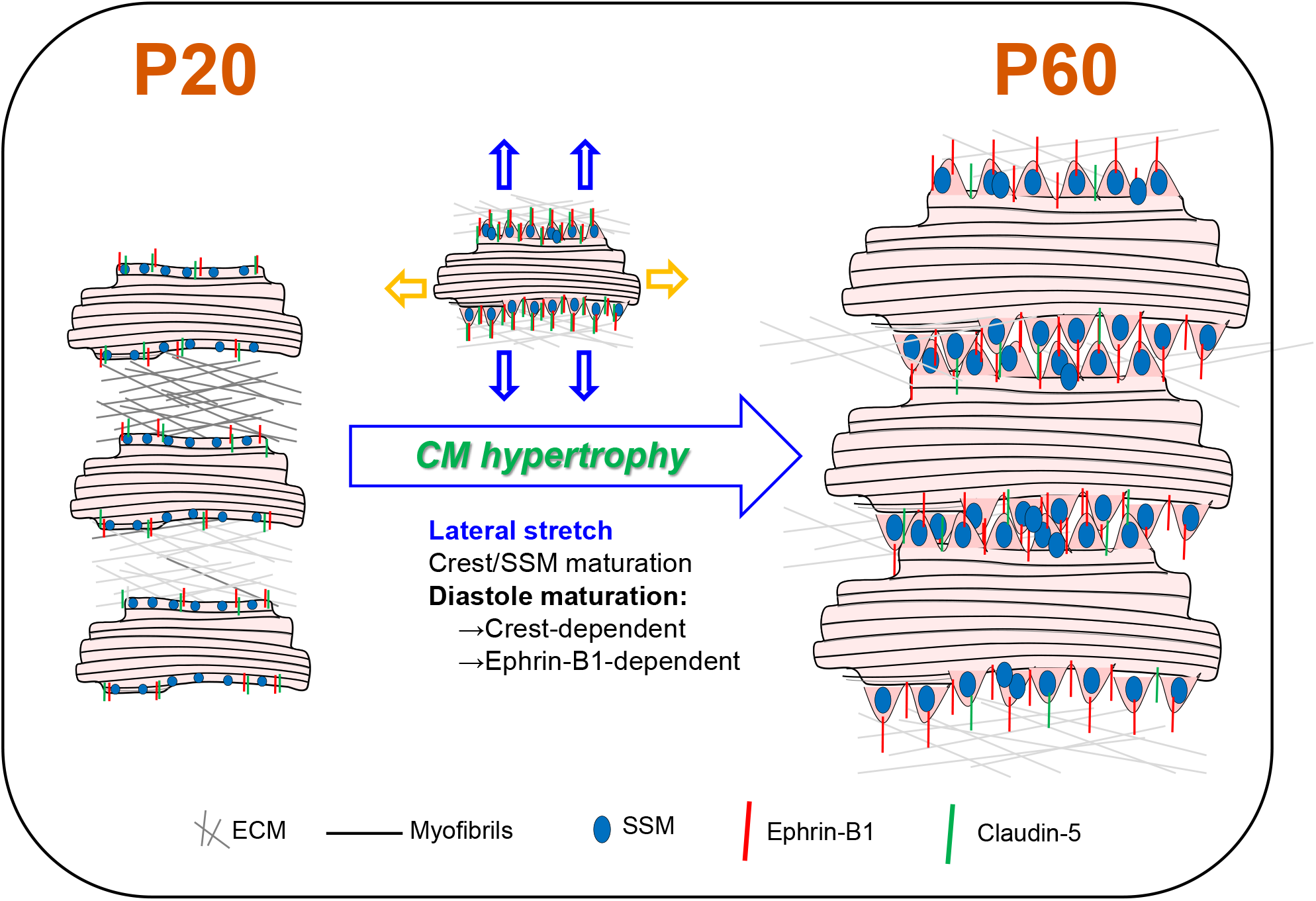
Schematic illustration of CM surface crest maturation between P20 and P60 supporting the setting of the adult diastolic function. At postnatal day 20 (P20), cardiomyocytes express low levels of Ephrin-B1 and its Claudin-5 partner at the lateral surface and surface crests/subsarcolemmal mitochondria (SSM) are still immature. CMs do not interact with neighbouring CMs at their lateral surfaces. After P20, CMs undergo substantial hypertrophy relying on elongation of both their long and short axes. From the short axis standpoint, the hypertrophy is independent from myofibril addition but relies on the maturation of the CM surface architecture. Hence, at P60, the increase in ephrin-B1/Claudin-5 expression at the lateral membrane promotes crest-SSM swelling but also crest-crest interactions between neighbouring CMs most likely leading to a lateral stretch of the CM. This Ephrin-B1-dependent lateral maturation of the CM surface allows maturation of the diastolic function.

Postnatal maturation of the heart in mammals has been paid much less attention than embryonic and fetal development. This would be a prerequisite for future pediatric clinical trials, which are seriously lacking. It also contributes to an significant knowledge gap that impedes research progression in regenerative medicine, since this postnatal maturation coincides with the proliferation blockage of the CM. Up to now, postnatal cardiac maturation has been essentially described until P20 in rodents, most likely related to the implementation of the typical adult rod shape of the CM. Very few studies have examined the transition from P20 to the adult stage, assuming that the genetic maturation of the heart is completed by P20 and that the heart merely undergoes a substantial growth beyond P20 until the young adult stage^14^. However, this period coincides with the weaning time in rodents (~P21) and thus with a critical nutritional/metabolic switch in the organism likely to influence heart function. So far, research has focused on the P0 (birth) to P20 postnatal window, during which the heart switches from a hyperplasic growth (CM proliferation) (~ P3-P5) to an hypertrophic growth (increase in CM size) (P20-21)^21–23^. Recent omics studies have detailed the molecular changes associated with the postnatal development of the mouse heart but only until P20^24,25^. Thus, the P0-P20 postnatal development period essentially relies on the ECM remodelling of the cardiac tissue and, at the CM level, myofibril and Ca^2+^ handling maturation, metabolic switch from glycolysis to major fatty acid oxidation and the transition from CM proliferation to CM hypertrophy^26,27^. This metabolic change most likely occurs as an adaptation of the postnatal heart to cardiac tissue oxygenation and the newborn diet, which essentially consists of mother’s milk and thus on high fatty acid availability. Our transcriptome analysis of mouse hearts reveals an additional developmental genetic program between P20 and P60 originating from all the cardiac cells of the left ventricle, thus indicating that the heart and the CM had not completed maturation by P20. Of interest, while the primary metabolic reliance of the heart on fatty acids and then glucose is already programmed between birth and P20, we found that the P20-P60 late postnatal stage programs the metabolic diversification of the heart. Such a new final metabolic adaptation of the heart coincides with the weaning process occurring around P20 in rodents, during which the newborn switches from milk to solid food^28^. This step is also in line with the capacity of the heart to metabolize a large panel of substrates to meet the energy demands of the adult stage^29^. Remarkably, we found that the *Bdh1* gene was increased by more than 37-fold between P20 and P60, most likely indicating that the capacity of the heart to oxidize KB (*coming essentially from the liver*) as a fuel is not only restricted to the adaptation of the failing heart, as previously shown^30^, but is also necessary for the adult physiological state. These results also agree with recent findings demonstrating that the human adult heart can use a large panel of substrates as a fuel, including a large proportion of KB^29^. Horton et al did not report a major cardiac phenotype under basal conditions in *Bdh1*^CM−/−^ mice^30^. However, more in-depth cardiac phenotyping of these mice would undoubtedly be necessary to depict the role of KB in the adult cardiac physiology, such as diastolic function.

An intriguing finding from our study is the physiological cardiac hypertrophy of the heart between P20 and P60, which, from a CM lateral standpoint, does not rely on a classical myofibril addition in the CM but, at least in part, on a ephrin-B1-dependent-i/ crest maturation through SSM swelling at the CM surface and -ii/ lateral stretching of the CM. During postnatal development, heart growth occurs primarily through CM proliferation (hyperplasia) but transitions rapidly after birth (~P5-7) to CM growth (hypertrophy)^22,23^ through new myofibril biogenesis. Although physiological CM hypertrophy has been described beyond P20 and is related to increased cardiac mass^14,31^, the underlying molecular mechanisms have never been explored. Piquerau *et al^14^* already reported and questioned such atypical hypertrophy of the rat heart after P21 without fiber addition, which was not classically related to the increase in heart-weight to body-weight index but conversely to a marked decrease. Here, we show that the P20-P60 hypertrophy relies on both an enlargement of the short and long axes of the CM. Interestingly, we demonstrated that the short axis elongation is dependent on the architectural maturation of the CM surface trough an ephrin-B1 mechanism. Thus, the ephrin-B1 lateral membrane protein, a partner and regulator of claudin-5^9^, participates in P20-P60 lateral CM hypertrophy, likely by bringing claudin-5 into the vicinity of neighboring CM, allowing intermittent but direct lateral crest-crest interactions (claudin-5 / claudin-5 interactions), crests/SSM maturation and the ensuing stretching of the CM lateral membrane (**Figure 8**). In agreement with this model, we previously showed that CMs from *Efnb1*^*CM−/−*^ KO mice exhibit substantially decreased levels of claudin-5 expression^9^, likely accounting for the lack of crest maturation and lateral crest-crest interactions in these adult KO mice. However, other mechanisms also likely contribute to the short-axis elongation, since it was only partially prevented by ephrin-B1 deletion. More specifically, lateral membrane interactions with the ECM, i.e. through integrin or the dystroglycan complex, that we previously showed to be independent of ephrin-B1^9^, could also play a role. In line with this assumption, our transcriptomic analysis identified the regulation of genes from the ECM remodelling pathway during the P20-P60 maturation period and the dystrobrevin-encoding gene (DTNA, **Figure X in the Supplemental Material**) from the dystroglycan complex. Apart from the short-axis, P20-P60 CM hypertrophy also depends on the elongation of the CM long-axis, which we show here to be independent from ephrin-B1 and which probably depends on the classical assembly of new sarcomeres at the myofibril extremities.

A new finding of our study is the specific maturation of SSM during the late P20-P60 postnatal period of the CM. Coinciding with this late cardiac mitochondria maturation in the CM, Piquerau et al^*14*^ previously reported a substantial increase in maximum respiratory capacity occurring after P21 and the adult stage in cardiac tissues from rats while the postnatal maturation of IFM occurred earlier. Biogenesis/maturation of cardiac mitochondria occur early during the embryogenesis, concurrently with the energy demand of the heart during the development period^16,32^. Thus, they can adapt their morphology and function according to the energy need and metabolic conditions of the cell^33,34^. In this field, most of the works on the postnatal period have focused on the characterization of the IFM (the most abundant in the adult CM) or the global cardiac mitochondria activity dedicated to the energy supply to the CMs for the contractile machinery, without distinguishing the different mitochondria subpopulations of the adult CM^35^. IFM maturation during the P0-P7 stage occurs both through swelling and through architectural reorganization along the myofibrils^14^, concomitantly with the well-known metabolic shift of the heart from glycolysis to a central oxidative metabolism more efficient for adult CM contraction^16^. It follows that IFM function is primarily dedicated to supplying the CM with the energy necessary for adult contraction. The delayed maturation of SSM at the CM surface compared to intracellular IFM supports the concept that these mitochondria regulate different CM functions. In agreement with this notion, we demonstrated in this study that crests/SSM specifically regulate heart diastole. Several results support this conclusion: i/ young adult *Efnb1*^*CM−/−*^ KO mice with immature SSM display diastolic defects with no impairment of systolic function, ii/ systolic function is already mature by P20 in rat hearts while the diastolic function is highly variable. Moreover, supporting our results about the late maturation of diastole, Zhou *et al* previously reported that the diastolic function in the left ventricle of mice matures around the weaning period^36^. In the future, it would be interesting to investigate whether cardiac SSM defects are a specific feature of diastolic dysfunction pathologies, such as HFpEF frequently associated with a metabolic syndrome^37^. In support of this concept, a recent study of myocardial gene expression signatures in human HFpEF reported that the ephrin-B1-encoding gene, that we demonstrated here as a specific determinant of the CM crests/SSM, is specifically downregulated in HFpEF heart patients compared with HFrEF^20^. Unfortunately, while several works have reported the influence of different cardiac pathologies on specific subpopulations of cardiac mitochondria^35^, these results require some caution, given the technical difficulty of specifically purifying and distinguishing the populations in the absence of reliable markers and due to the fact that these mitochondria are morphologically highly similar^12^. Today, electron microscopy still remains the gold standard for accurately examining the cardiac mitochondria subpopulations.

An essential and new finding of our study is the identification of the ephrin-B1/crest/SSM module at the lateral membrane of the CM as a specific determinant of the physiological diastolic function of the adult heart. So far, the diastolic determinants have been proposed only in the context of diastolic dysfunction in cardiac pathologies such as HFpEF^38^. However, until rather recently, the lack of specific and compelling therapeutics in HFpEF underlines our limited knowledge of the control of diastole. This is overcomplicated in the context of HFpEF due to the contributions of several comorbidities delineating highly heterogeneous clinical features^38^. These last months, the discovery of empaglifozine, a iSGLT2 antidiabetic drug, as the first effective therapeutics for HFpEF patients^39^, questioned the molecular mechanisms underlying the cardiac benefit, given the lack of SGTL2 expression in the heart. However, a beneficial and indirect role through metabolism regulation has been postulated^40^. It follows that the control of the diastole could arise, in part, from a systemic metabolic regulation. Thus, the extent to which iSGTL2 could regulate the crest/SSM module at the CM surface in a pathological context should be consider in the future. Here, our study demonstrates that *Efnb1* deletion specifically in the CM recapitulates at the young adult stage (2 months) some features of the diastolic dysfunction depicted in HFpEF, i.e. an elevated EDP and altered filling patterns combined with exercise intolerance. One important concern is understanding how ephrin-B1 at the lateral membrane of the adult CM can impact diastolic function. We previously demonstrated that ephrin-B1 controls the architecture of the lateral membrane and the overall adult rod shape of the CM^9^. Here, we now further demonstrated that ephrin-B1 controls the maturity of the crests/SSM architectural motif at the lateral membrane, thus playing a key role in the adult crest-crest interactions between neighboring CMs and in overall tissue cohesion. These lateral CM interactions likely dictate a mechanical lateral stretch of the CM, allowing perfect stacking between myofibril layers. Consequently, crest-crest interactions might contribute to controlling the relaxation length of the sarcomere, a mechanism that we previously suggested depends on crest height^12^. This ephrin-B1-dependent lateral stretch of the CM is also necessary for the spatial arrangement of the T-tubules and most likely its ensuing function during the P20-P60 development, as supported by the T-tubule disorganization in the *Efnb1*^*CM−/−*^ KO mice. Thus, by altering the T-tubule structure, lack of ephrin-B1 could influence the Ca^2+^ entry / Ca^2+^ exit. It is worth noting that T-tubule disorganization in the CMs of *Efnb1*^*CM−/−*^ KO mice is progressive with mild disorganization correlating with diastolic dysfunction only, while extensive disorganization is observed in HFrEF. Although defects in T-tubule architecture/function are a hallmark of HFrEF^41^, it has been poorly examined in HFpEF, only one recent study reported that this mechanism is etiology-dependent^42^. An exciting feature of the *Efnb1*^*CM−/−*^ KO mice is the progressive cardiac phenotype starting from a primarily diastolic defect that progressively switches toward a mild and then severe HFrEF and finally to death, thus highlighting the substantial cardioprotective role of the surface crests/SSM of the lateral membrane. Although this switch toward HFrEF is not a common feature of HFpEF patients^43^, definitive conclusions cannot be reached on the natural evolution of HFpEF, given that HFpEF patients received medication for their different comorbidities, which could protect them from HFrEF. In the future, an accurate characterization of the CM surface crests/SSM in different HFpEF models with diastolic dysfunction will undoubtedly shed light on whether crest disruption is a hallmark of HFpEF and could thus contribute to the setting of the pathology.

## NONSTANDARD ABBREVIATIONS AND ACRONYMS

AFM: Atomic force microscopy
CM: Cardiomyocyte
DBP: Diastolic blood pressure
ECM: Extracellular matrix
EDP: End diastolic pressure
FA: Fatty acid
GO: Gene ontology
HF: Heart Failure
HFrEF: Heart Failure with reduced ejection fraction
HFpEF: Heart Failure with preserved ejection fraction
ID: Intercalated disk
IFM: Interfibrillar mitochondria
IVRT: Isovolumic relaxation time
KB: Keytone bodies
KO: Knock-out
LVPWd: left ventricular posterior wall in diastole
LVEDV: Left ventricular end-diastolic volume
LVEDD: Left ventricular end-diastolic diameter
LVIDd: left ventricular internal diameter in diastole
LV-GLS: left ventricular global longitudinal strain
P5: Postnatal day 5
SBP: Systolic blood pressure
SSM: Subsarcolemmal mitochondria
SCIM: Scanning ion-conductance microscopy
TEM: Transmission electron microscopy
WT: wild-type

## Acknowledgements

We thank Claire Naylies and Yannick Lippi (Toxalim, Université de Toulouse, INRAE, ENVT, INP-Purpan, UPS, Toulouse, France) for their contribution to microarray fingerprints acquisition and microarray data analysis carried out at GeT Genopole Toulouse Midi-Pyrénées facility (https://doi.org/10.15454/1.5572370921303193E12).

We are also grateful to TRI Genotoul network facilities, specifically ANEXPLO platform (Toulouse) for help with echocardiography, the “Centre de Microscopie Electronique Appliquée à la biologie-CMEAB» (Faculté de Médecine Rangueil-Toulouse) and the Cellular Imaging Facility-I2MC (Toulouse).

