## Supplementary Methods and Figures for "Crest maturation at the cardiomyocyte surface contributes to a new late postnatal development stage that controls the diastolic function of the adult heart"

### Supplementary Materials

### SUPPLEMENTARY METHODS

#### Animal models

CM-specific (cKO) *efnb1* KO mice (*Efnb1*$^{CM-/-}$KO) have already been described[1]. *Efnb1* WT and cKO mice were kept in a mixed S129/S4 × C57BL/6 background. All studies were performed on male and age-matched mice. Studies on rats were performed on male Sprague-Dawley rats (purchased from Envigo, Huntingdon, United-Kingdom). Experimental animal protocols were carried out in accordance with the French regulation guidelines for animal experimentation and were approved by the French CEEA-122 ethical committee.

#### Transmission electron microscopy (TEM) and quantitative analysis

Heart samples were specifically processed to preserve CM surface crests as previously described[2]. Thus, immediately after removal, the heart was washed in cold PBS for several seconds, then cautiously sliced (~1 $mm^3$ biopsies) on a glass slide lying on an ice bed and fixed in 2% glutaraldehyde in Sorensen's buffer at 4°C and processed for TEM as previously described[3].

The quantification of crest heights and sarcomere lengths/heights was performed on TEM micrographs as previously described[2]. The crest heights were measured from the top of the crest to the first myofibril top layer using FiJi software and the SSM number was quantified for each crest. The sarcomere lengths were measured similarly on the first myofibril layer below the CM surface crests. Myofibril numbers were quantified on the longitudinal axis of the CM at the level of their anchorage on an entire intercalated disk from top to bottom of the cell.

#### Echocardiography, speckle tracking echocardiography / strain analysis and Doppler imaging

All echocardiographies were performed under isoflurane anesthesia (1.5% - 2%). The animals' limbs were taped over the metal ECG leads. Body temperatures were carefully monitored and maintained as close to 37°C as possible during the entire procedure. For **Figure 2C (**P20 and P60 rats) transthoracic echocardiography was carried out using the Vevo 3100 (Fujifilm VisualSonics), equipped with a 21 MHz transducer (MX250). For **Figure 4A** (P20 and P60 rats), transthoracic echocardiography was carried out using the Acuson NX3 Elite (Siemens Healthineers) equipped with a high-frequency 16-MHz linear transducer (VF16-5 probe, Siemens Healthineers). For **Figure 6A** (adult mice), transthoracic echocardiography was carried out using a Vevo 2100 (Fujifilm VisualSonics), equipped with a 25 MHz transducer (MS250).

Left ventricular (LV) parasternal long-axis and short-axis 2D view in M-mode was performed at the papillary muscle to assess LV wall thicknesses and internal diameters. Left ventricular ejection fraction (LVEF) was also calculated (%) from a B-mode parasternal long-axis view by tracing endocardial end-diastolic and end-systolic borders to estimate LV volumes (EF % = [end-diastolic volume (EDV)-end-systolic volume]/EDV×100).

The global longitudinal strain was measured from parasternal long-axis images using speckle-tracking-based imaging to evaluate global cardiac performance. Diastolic function was assessed by pulse-wave Doppler imaging of the mitral flow obtained from the apical four-chamber view. Early (E) and late (A) peak transmitral flow velocity and isovolumic relaxation time (IVRT) were measured, and the E/A ratio was calculated. Longitudinal motion of the mitral annulus was captured by tissue Doppler imaging. Mitral annular velocities (E′ and A′ peaks) were measured and the E/A ratio and E/E' were calculated. Left atrium dimensions (anteroposterior diameter) were compared to the aortic diameter in the parasternal long axis. All measurements were quantified and averaged for three cardiac cycles and obtained by an examiner blinded to the genotype of the animals.

**Hemodynamic measurements of left ventricular pressure**

Catheterization was performed before sacrifice, under isoflurane anesthesia (2%). The animals were monitored with ECG leads and body temperatures were carefully monitored and maintained as close to 37°C as possible during the entire procedure. Hemodynamic parameters were measured using a Science pressure catheter (Transonic) connected to the PowerLab 8/35 data acquisition system (AD Instruments). The catheter was inserted into the right carotid artery, and after stabilization for 5 min, arterial blood pressure was recorded. After that, the catheter was advanced into the ascending aorta and then into the left ventricle under pressure control. After stabilization for 5 min, signals were continuously recorded. Trace analysis was then performed using the LabChart Pro software system (AD Instruments) to determine systolic and diastolic blood pressure, LV end-diastolic pressure (LVEDP), the maximum slope of the LV systolic pressure increment (dP/dtmax) and diastolic pressure decrement (dP/dtmin), and the relaxation time constant (Tau) was calculated from the LV pressure trace.

**Exercise exhaustion test**

The maximum stress test with measurement of gas exchange was carried out according to the method described by Kemi *et al.*[4] using the TSE CaloTreadmill system, a fully computerized, electronically controlled system to investigate exercise calorimetry in mice and rats. CaloTreadmills are enclosed by an air-tight transparent housing that can be connected to a fully automated indirect gas calorimetry system operated by PhenoMaster software.

After two days of acclimatization to the treadmill, the stress test was carried out in mice from the two experimental groups, respecting heterogeneity between the two populations on each run. The mice warmed-up on an inclined treadmill (25°) at a speed of 5 $cm.s^{-1}$ for 5 min. Then, the mice ran at 15 $cm.s^{-1}$ for 5 min, and the speed was increased every 5 minutes by 5 $cm.s^{-1}$ until reaching 25 $cm.s^{-1}$. Finally, the speed was increased by 3 $cm.s^{-1}$ every 2 minutes until exhaustion. Exhaustion was defined as the inability of the mouse to return to running within 10 s of direct contact with an electric-stimulus grid. Calorimetric parameters, such as oxygen consumption ($VO_2$), carbon dioxide production ($VCO_2$), respiratory exchange rate (RQ) and energy expenditure (EE) are determined at fixed temporal intervals while the test animal is moving according to an experimenter-defined activity profile.

**Heart sections**

For paraffin-embedded heart sections, hearts were excised and immediately fixed in 10% formalin (24 h), 4% formalin (48 h) and then embedded in paraffin and sectioned (transversal sections) at 5 μm intervals. For frozen cardiac tissue sections, hearts were embedded in OCT, frozen in liquid nitrogen and 5 μm frozen sections were fixed with ice-cold acetone. as previously described[1].

**Immunohistochemistry and confocal imaging**

Formalin-fixed paraffin-embedded or frozen cardiac tissue sections were used for immunohistochemical analysis. For paraffin-embedded sections, samples were deparaffinized, rehydrated and subjected or not to heat-induced epitope retrieval depending on the antibodies. For cryosections, samples were permeabilized (0.5% Triton X100-PBS), blocked (Dako Protein Block, Dako, X0909 or 10% normal goat serum, Dako, X0907 or 0.5% BSA/PBS), and stained overnight at 4°C with the primary antibodies, then 1 h RT with the fluorescent secondary antibodies all diluted in 0.1% BSA / 0.1% Tween 20 / 0.5% Triton X-100-PBS. Nuclei were visualized with DAPI (Sigma Aldrich, 32670). Cell membranes were stained with OG488-conjugated wheat germ agglutinin (WGA, Life Technologies, W21404). All images were acquired on Zeiss LSM 780 confocal microscope using Zen 2011 software (Carl Zeiss).

**Antibodies**

The primary antibodies in this study were as follows: goat anti-ephrin-B1 (R&D); mouse anti-α sarcomeric actinin (Sigma); mouse anti-connexin 43 (Millipore); rabbit anti-N-cadherin (Epitomics); rabbit anti-desmoplakin 1/2 (ARP); rabbit anti-claudin-5 (Acris antibodies); mouse anti-GAPDH (Abcam), mouse anti-ryr (Abcam), mouse anti-caveolin-3 (BD Transduction Laboratories$^{TM}$). The secondary fluorescent antibodies used in this study were as follows: donkey anti-goat OG-488 Alexa Fluor; goat anti-rabbit OG-488-Alexa Fluor, donkey anti-mouse OG-488 Alexa Fluor, donkey anti-goat Cy3-Alexa Fluor and donkey anti-rat Cy3-Alexa Fluor all obtained from Invitrogen.

**2D quantification of CM area/density**

For *in situ* quantification of CM surface area and density, deparaffinized heart slides were stained with OG488-conjugated-WGA and CM area and density were measured in transversal heart cross-sections by manually tracing the cell contour on images of whole hearts acquired on a digital *NanoZoomer* (Hamamatsu) slide scanner using Zen 2011 software (Carl Zeiss). The experimenter was blinded to groups and genotypes.

**Quantification of cardiac fibrosis**

Paraffin-embedded heart sections were stained using the Trichrome stain kit (Connective Tissue Stain, Abcam, ab150686). Fibrosis was quantified on Masson's trichrome-stained heart sections using an ImageJmacro from images of whole heart sections acquired on a digital *NanoZoomer* (Hamamatsu) slide scanner. The experimenter was blinded to the mouse genotype.

**Quantification of T-Tubules (TT) regularity in the cardiac tissue.** TT architecture was quantified in LV myocardium from 4% PFA fixed-rat cardiac biopsies, and processed for Caveolin-3 fluorescent immunostaining. Caveolin-3 was chosen as an indirect TT marker as it was previously shown that caveolin-3 and TT markers co-localized intracellularly at the Z-disk in adult CMs[5]. Images from caveolin-3-stained cardiac biospsies were acquired with a laser-scanning confocal microscope (Carl Zeiss LSM900), with 63x/1.4 oil objective and a 100 nm sampling. From these images, several regions of interest were extracted, each corresponding to different cells. This extraction was performed on ImageJ by manually rotating the image, to have the CMs long axis oriented along the image horizontal axis. Then regions of interest with a fixed size of 500 by 35 pixels (corresponding to an area of 50 µm by 3.5µm) were manually drawn and duplicated from the image. On these regions, the quantitative TT power measurement was performed with a Python script, written in a Jupyter Notebook. At first a two-dimensional Fast Fourier Transformation (FFT2D) is performed. From the FFT2D the power spectrum P_2D is calculated as: P_2D= | 〖FFT〗 _2D |^2/N with N the total number of pixels. From the power spectrum image, a one-dimensional power spectrum P_1D is retrieved by extracting the middle horizontal line. Finally, TT power (indicative of the regular organization of TT system) was measured on P_1D as the amplitude of a Gaussian curve fitted on a peak located between 0,45 $\mu m^{-1}$ and 0,7 $\mu m^{-1}$ and frequency (indicative of  as the center of the Gaussian curve. The script uses the libraries Numpy, Pandas, AICSImageIO, SciPy and Matplotlib.

**Cardiomyocyte isolation and contractility**

Unloaded cell shortening and calcium transients were measured in freshly isolated ventricular myocytes, prepared as described previously[6]. Isolated cardiomyocytes were loaded with Indo-1 AM (10 µM Invitrogen, France) at room temperature for 30 min and then washed out with the free HEPES-buffered solution (117 mM NaCl, 5.7 mM KCl, 4.4 mM $NaHCO_3$, 1.5 mM $KH_2PO_4$, 1.7 mM $MgCl_2$, 21 mM HEPES, 11 mM glucose, 20 mM taurine, pH 7.2) containing

1.8 mM $Ca^{2+}$ [7]. Unloaded cell shortening and calcium concentration [$Ca^{2+}$] (indo 1 dye) were studied using field stimulation (1 Hz, 22°C, 1.8 mM external $Ca^{2+}$). Sarcomere length (SL) and fluorescence (405 and 480 nm) were simultaneously recorded (IonOptix System, Hilton, USA).

**Western blotting**

For protein analysis from total cardiac extracts, cardiac tissue samples were directly lysed in RIPA buffer (25 mM Tris-HCl pH 7.6, 150 mM NaCl, 1% NP-40, 1% NaDOC, 0.1% SDS) in the presence of a protease inhibitor cocktail (Roche). The protein concentration of extracts was determined by the Lowry method (Bio-Rad) and equal amounts of proteins were subjected (50 µg) to SDS-PAGE and transferred to nitrocellulose membranes (Millipore). Proteins were detected with primary antibodies followed by horse radish peroxidase-conjugated secondary antibodies to mouse or rabbit IgG (GE Healthcare) using an enhanced chemiluminescence detection reagent (GE Healthcare). Protein quantification was obtained by densitometric analysis using ImageQuant 5.2 software and was normalized to that of GAPDH expression and expressed in arbitrary units (A.U.).

**RNA extraction and microarray gene expression studies**

RNA was extracted from left ventricle heart samples using the Tri-reagent method[8] (Sigma Aldrich France). Gene expression profiles were performed at the GeT-TRiX facility (GénoToul, Génopole Toulouse Midi-Pyrénées) using Agilent Sureprint G3 Mouse GE v2 microarrays (8x60K, design 074809) following the manufacturer's instructions. For each sample, cyanine-3 (Cy3)-labeled cRNA was prepared from 200 ng of total RNA using the One-Color Quick Amp labeling kit (Agilent Technologies) according to the manufacturer's instructions, followed by Agencourt RNAClean XP (Agencourt Bioscience Corporation, Beverly, Massachusetts). Dye incorporation and cRNA yield were checked using the Dropsense™ 96 UV/VIS droplet reader (Trinean, Belgium). 600 ng of Cy3-labelled cRNA were hybridized on the microarray slides following the manufacturer's instructions. Immediately after washing, the slides were scanned on Agilent G2505C Microarray Scanner using the Agilent Scan Control A.8.5.1 software and fluorescence signal extracted using Agilent Feature Extraction software v10.10.1.1 with default parameters.

Microarray data and experimental details are available in NCBI's Gene Expression Omnibus[9] and are accessible through GEO Series accession number GSE196257 (https://www.ncbi.nlm.nih.gov/geo/query/acc.cgi?acc=GSE196257 ).

**Microarray data statistical analysis**

Microarray data were analyzed using R (R Core Team, 2018) and Bioconductor packages[10] as described in GEO accession GSE196257. Raw data (median signal intensity) were filtered, log2 transformed and normalized using the quantile method[11].

A model was fitted using the limma lmFit function[12]. Pair-wise comparisons between biological conditions were applied using specific contrasts. A correction for multiple testing was applied using the Benjamini-Hochberg procedure[13] to control the false discovery rate (FDR). Probes with FDR $\leq 0.05$ were considered to be differentially expressed between conditions.

Hierarchical clustering was applied to the samples and the differentially expressed probes using the Pearson correlation coefficient as distance and Ward's criterion for agglomeration. The clustering results are illustrated as a heatmap of expression signals. The enrichment of gene ontology (GO) biological processes and KEGG pathways was evaluated using the enrichGO function and enrichKEGG functions from the clusterProfiler R package[14] .

##### Gene clustering according to the cardiac cell populations

Processed single-cell RNA-seq data from Tucker *et al.*[15] were retrieved from the Broad Institute's Single Cell Portal (study ID SCP498). Highly variable genes were selected on the scaled logNormalize dataset to perform cell clustering using Seurat 4.0.1 (using the default parameters in the FindVariableFeatures function in the Seurat R package). The uniform manifold approximation and projection (UMAP) were performed based on the first 30 principal components. Cell identities were made available by Tucker *et al.*[15] and used to color the cells on the UMAP plot.

##### Data analysis-Statistics

The n number for each experiment and analysis is stated in each figure legend. The normality was tested using the Shapiro-Wilk normality test. Results are reported as mean ± SD. An unpaired Student t-test for parametric variables was used to compare two groups. One-way ANOVA with Tukey post-hoc test was used for multiple group comparisons (> 2 groups). Two-way ANOVA with Tukey post-hoc test was used for multiple group comparisons with two independent variable such as age (P20 and P60) and genotype (WT and cKO). The level of significance was assigned to statistics in accordance with their p values: $p \leq 0.05$ *; $p \leq 0.01$ **; $p \leq 0.001$ ***; $p \leq 0.0001$ ****). All graphs were generated using v9.2.0 (GraphPad Inc, San Diego, California).

#### Method references

11. Bolstad, B.M., Irizarry, R.A., Astrand, M. & Speed, T.P. A comparison of normalization methods for high density oligonucleotide array data based on variance and bias. *Bioinformatics* **19**, 185-93 (2003).
12. Ritchie, M.E. et al. limma powers differential expression analyses for RNA-sequencing and microarray studies. *Nucleic Acids Res* **43**, e47 (2015).
13. Benjamini, Y. Controlling the false discovery rate: a practical and powerful approach to multiple testing. *London, ROYAUME-UNI, Royal Statistical Society* (1995).
14. Yu, G., Wang, L.G., Han, Y. & He, Q.Y. clusterProfiler: an R package for comparing biological themes among gene clusters. *OMICS* **16**, 284-7 (2012).
15. Tucker, N.R. et al. Transcriptional and Cellular Diversity of the Human Heart. *Circulation* **142**, 466-482 (2020).

#### Data Supplement-Figure Legends

**Figure I. Ultrastructure of the neonatal CM at birth.** TEM micrographs showing representative CM within the cardiac tissue at postnatal day 0 (birth) with plasma membrane protrusions (arrows) between two Z-lines.

**Figure II. Interfibrillar mitochondria (IFM) during postnatal maturation.** TEM micrographs showing representative IFM (stars) during rat postnatal maturation at postnatal day 0 (P0), 5 (P5), 20 (P20) and 60 (P60).

**Figure III**. **Maturation of CM morphology during the postnatal period.** Confocal immunofluorescent staining of cell membranes (wheat germ agglutinin-OG488 (WGA), green) and CMs ($\alpha$-actinin, red) performed on paraffin-embedded longitudinal heart sections from rats of varying postnatal ages (P0, P3, P5, P10, P20, P60).

**Figure IV. Organization of CM myofibrills during the postnatal period.** Confocal immunofluorescent staining of CM sarcomere Z-lines ($\alpha$-actinin, red) and CMs cell surface (wheat germ agglutinin-OG488 (WGA), green) performed on PFA-fixed Fresh biopsies from left ventricles of rats from varying postnatal ages (P20, P60).

**Figure V**. **Neighboring CMs establish direct lateral physical contacts through crest-crest interactions only at the adult stage.** Illustrative TEM micrographs of P60 rat LV myocardium showing intermittent tight junctions (arrows) between the lateral surface crests of two adjacent CMs.

**Figure VI. Maturation of the intercalated disk of CMs during the postnatal period.** Representative confocal immunofluorescent staining/imaging of N-cadherin (*upper panels*), desmoplakin *(middle panels*) and connexin-43 (*lower panels*) performed in heart transverse cryo-sections from P0, P5, P20 and P60 rats. DAPI staining=cell nuclei.

**Figure VII. Organization of the T-tubule network during the late postnatal period.** Representative confocal immunofluorescent staining/imaging of ryanodine receptor (RyR) (*upper panels*) and Caveolin-3 (*middle panels*), as T-Tubule (TT) markers, in isolated CMs (**A**) or LV myocardial tissue (**B**) and showing a spatial misalignment (arrows) of these proteins in P20 rats compare to CMs from P60 rats. (**C**) Quantification of the TT regularity (along the periodic striation on sarcomere Z-lines) and frequency (periodicity on Z-lines) performed on LV myocardial tissue from P20 or P60 rat hearts (P20, n=4 rats; P60 n= 2 rats; ~5-20 CMs / rat).

**Figure VIII. Kinetics of CM hypertrophy during the late postnatal period.** CM area quantified from wheat-germ agglutinin (WGA)-stained heart cross-sections (transverse) obtained from P20, P30, P45 and P60 old rats (P20 n=4, P30 n=3, P45 n=4, P60 n=4 rats ~ 400 CMs/rat). Data are presented as mean ± SD; one-way ANOVA with Tukey post-hoc test for four group comparisons * $P < 0.05$, *** $P < 0.001$, ns, not significant.

**Figure IX. Late postnatal growth of the morphology of isolated CMs.** (*upper panels*) Representative light microscopy photographs of isolated CMs from P20 and P60 rat hearts. (*lower panel*) CM long and short axis lengths quantified on CMs isolated from P20- or P60-old rat hearts (~ 200 CMs/rat; n = 3-4 per group). Data are presented as mean ± SEM Unpaired Student's *t*-test for two group comparisons *** $P < 0.001$.

**Figure X. Hypertrophy of CMs in mice during the late postnatal period.** (**A**) CM area quantified from wheat-germ agglutinin (WGA)-stained heart cross-sections (transverse) obtained from P20 and P60 mice (~ 450 CMs/mouse; n=5 per group); (**B**) CM long and short axes quantified from WGA-stained heart cross-sections (myocardium) obtained from P20 and P60 mice (~ 120 CMs/mouse; n=3 per group). Data are presented as mean ± SD. Unpaired Student's *t*-test for two group comparisons **** $P < 0.0001$.

**Figure XI. Late postnatal maturation of the myofibrils.** Illustrative transversal TEM micrographs of rat LV myocardium showing larger distances between the thick myosin filaments at P60 than at P20.

**Figure XII. Signaling pathways upregulated or downregulated during the late postnatal period in left ventricles from WT mice.** Full gene ontology (GO) analysis of upregulated or downregulated genes between P60 and P20 in mouse left ventricles. The false discovery rate is provided for each category.

**Figure XIII. Regulation of cardiomyocyte and metabolic markers during the late postnatal period.** Expression level of markers of cardiomyocytes (**A**) or metabolic-pathways (**B**) in the different cardiac cell populations are plotted onto the UMAP plot. Color key from gray to blue indicates relative expression level from low to high.

**Figure XIV. *Efnb1*-specific knockdown in the CM impairs the lateral stretch of the CM.** (**A**) Quantification of sarcomere heights from TEM micrographs obtained from P20 or P60 *Efnb1*$^{CM-/-}$ KO and WT mouse hearts (P20 WT n=4, P20 cKO n=4, P60 WT n=5; P60 cKO n=6; 4 to 8 CMs/mouse, ~ 50 sarcomeres/mouse). (**B**) (*left panels*) Quantification of the lateral membrane interspace between two neighboring CMs from TEM micrographs obtained from P20 or P60 *Efnb1*$^{CM-/-}$ KO and WT mouse hearts (P20 WT n=1, P20 cKO n=2, P60 WT n=3; P60 cKO n=3; ~ 40 lateral membrane spaces/mouse) and (*right panel*) representative TEM. Data are presented as mean ± SD. Two-way ANOVA with Tukey post-hoc test for four group comparisons ** $P<0.01$, ns, not significant.

### Data Supplement

*Figure I - Karsenty et al.*

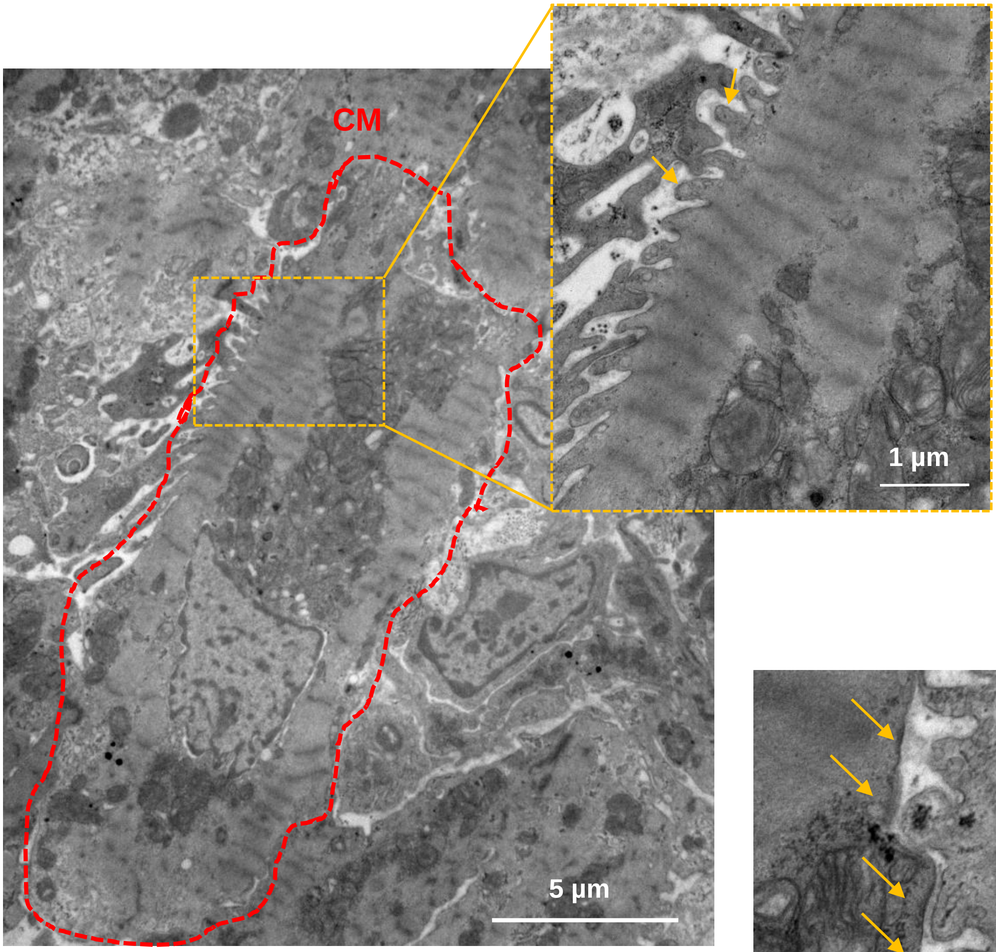


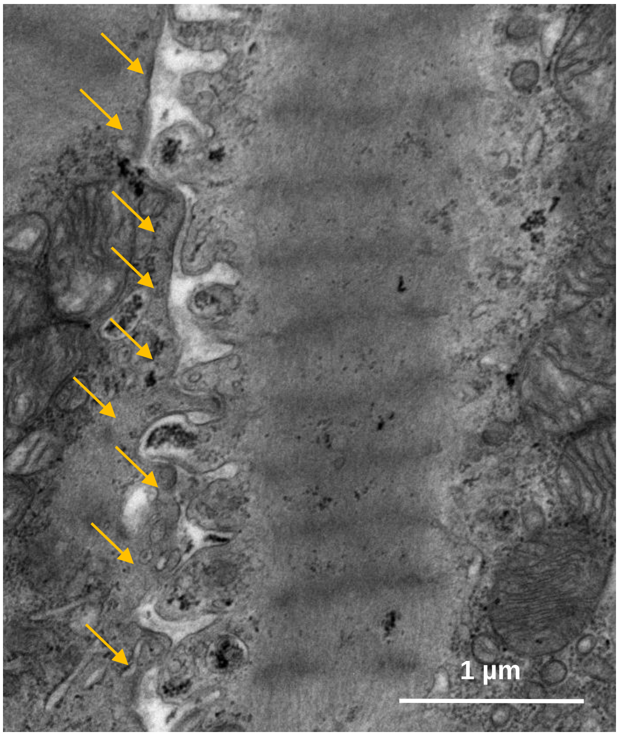


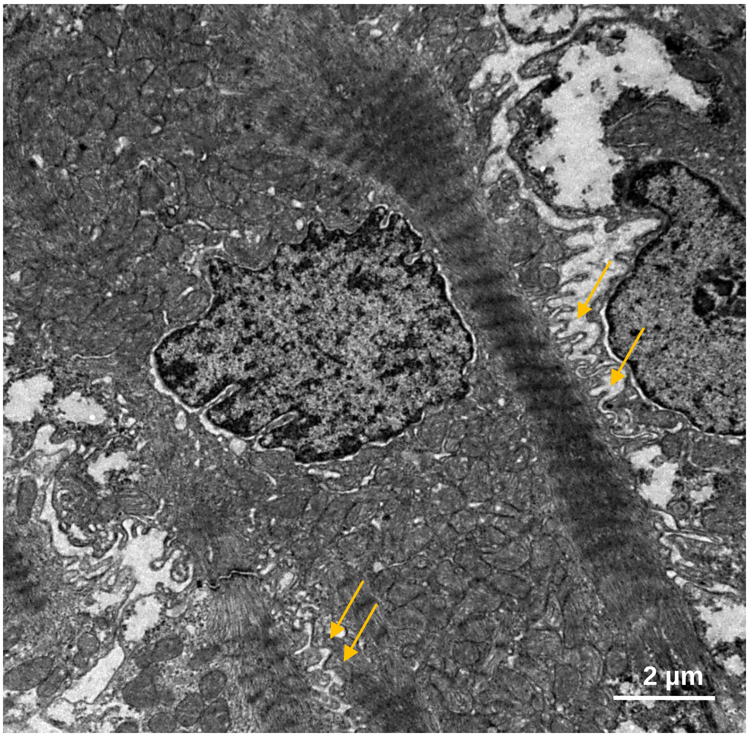

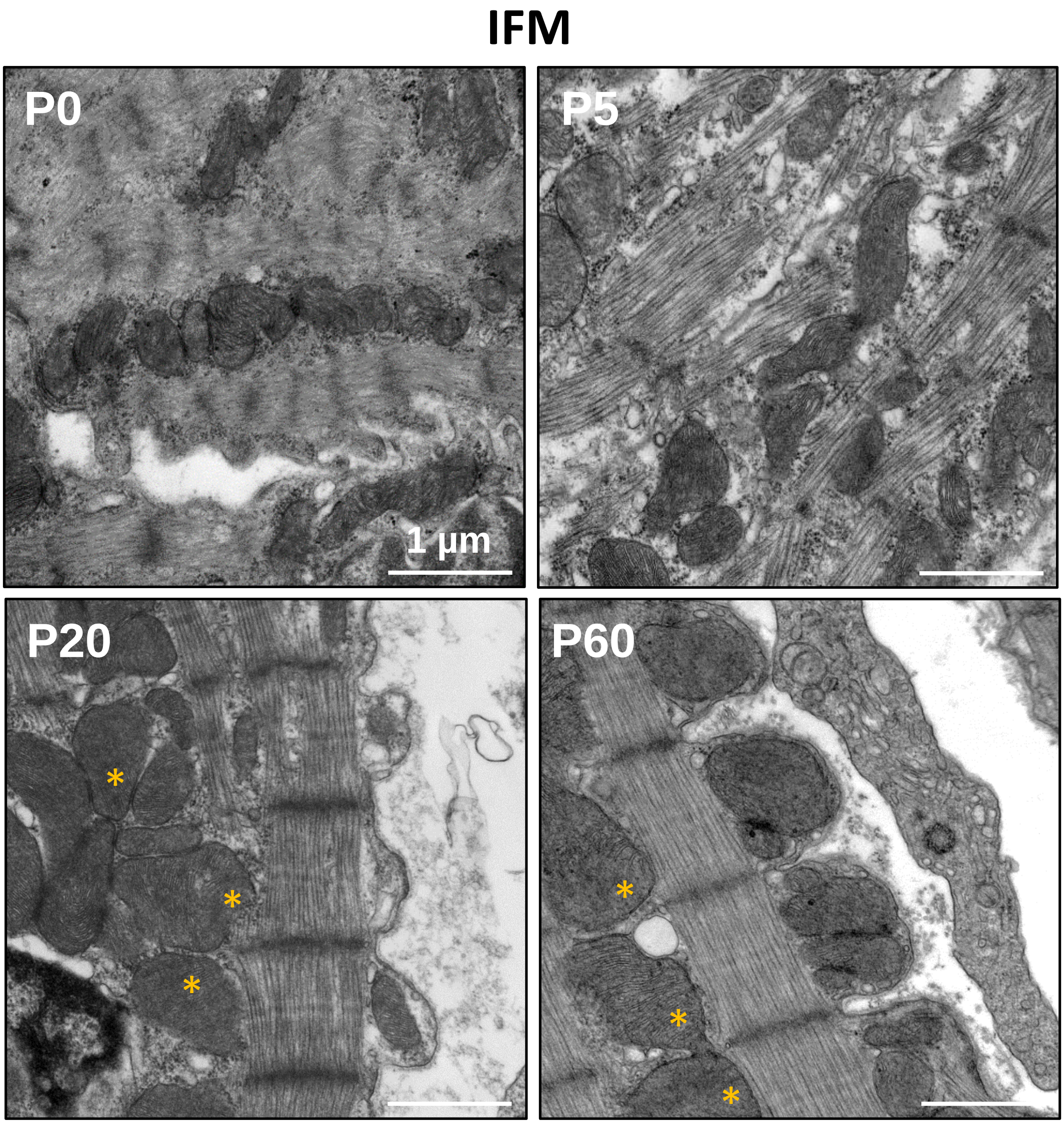
IFM
P0
1 µm
P5
P20
P60

*Figure III - Karsenty et al.*

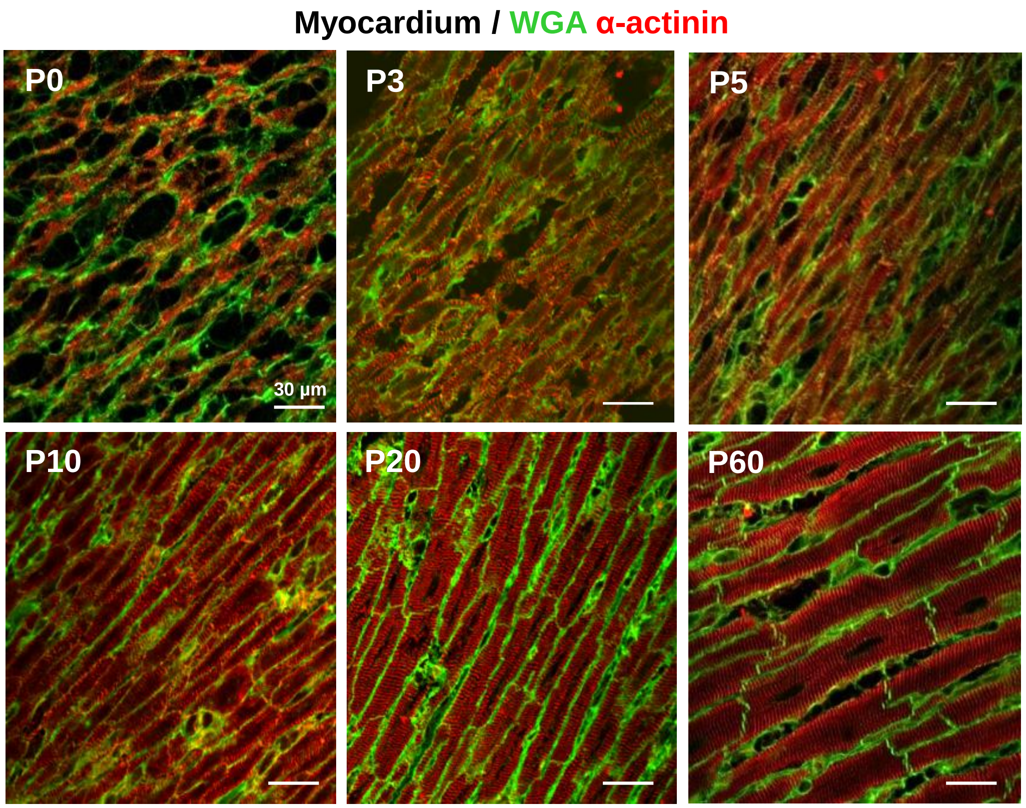


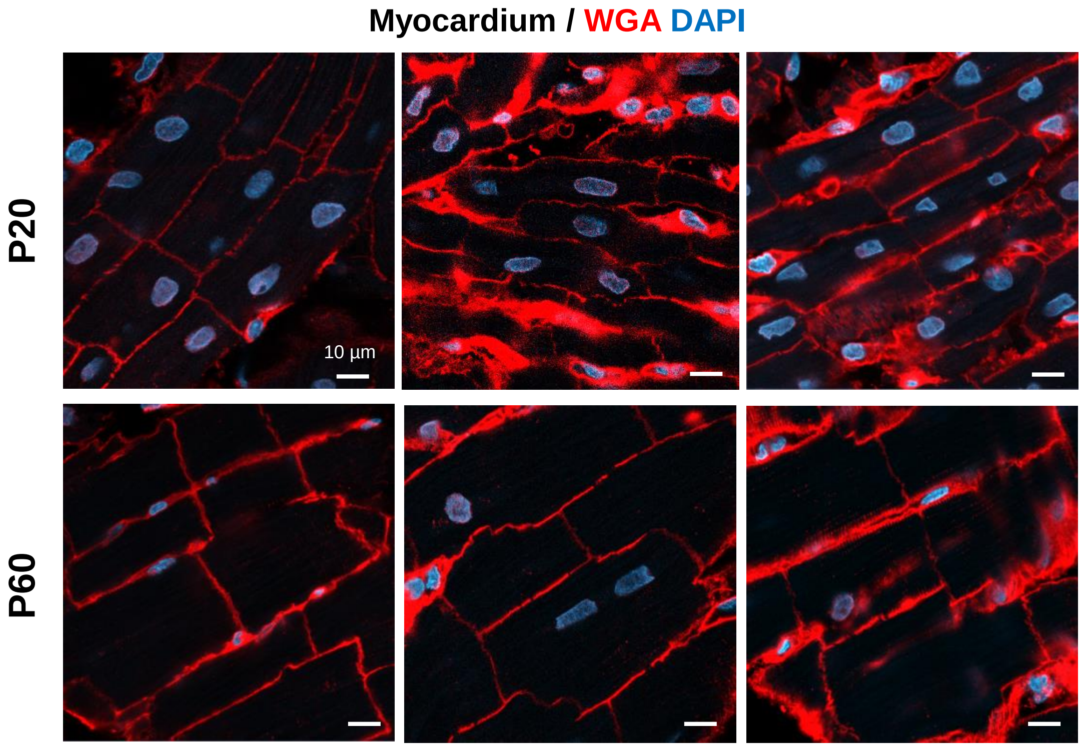

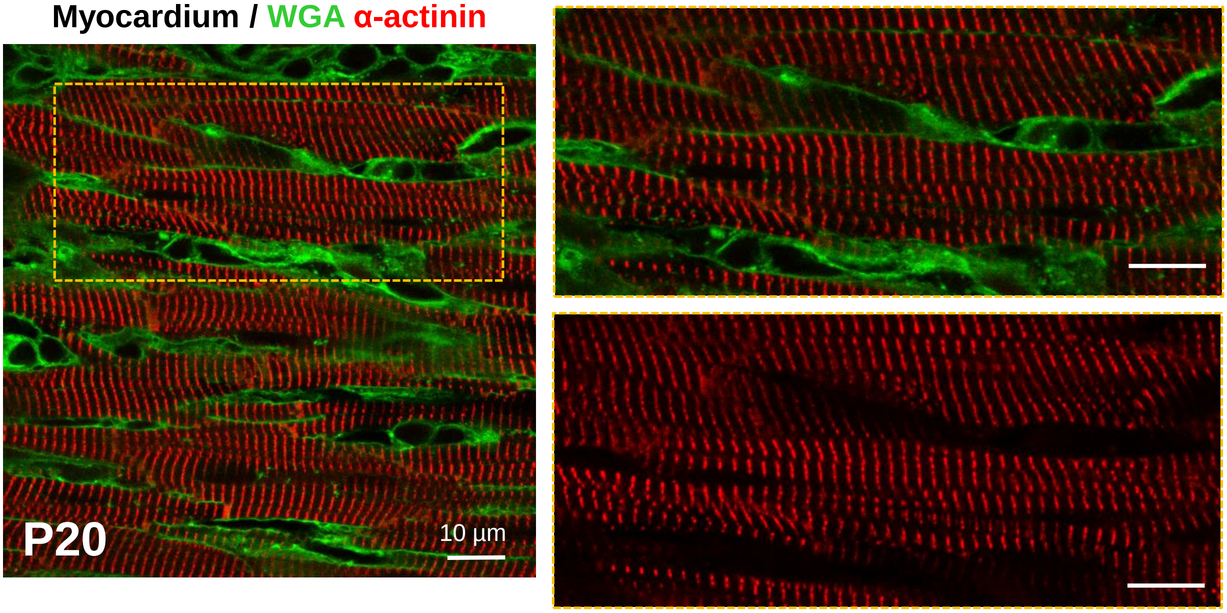
Myocardium / WGA α-actinin
P20
10 µm

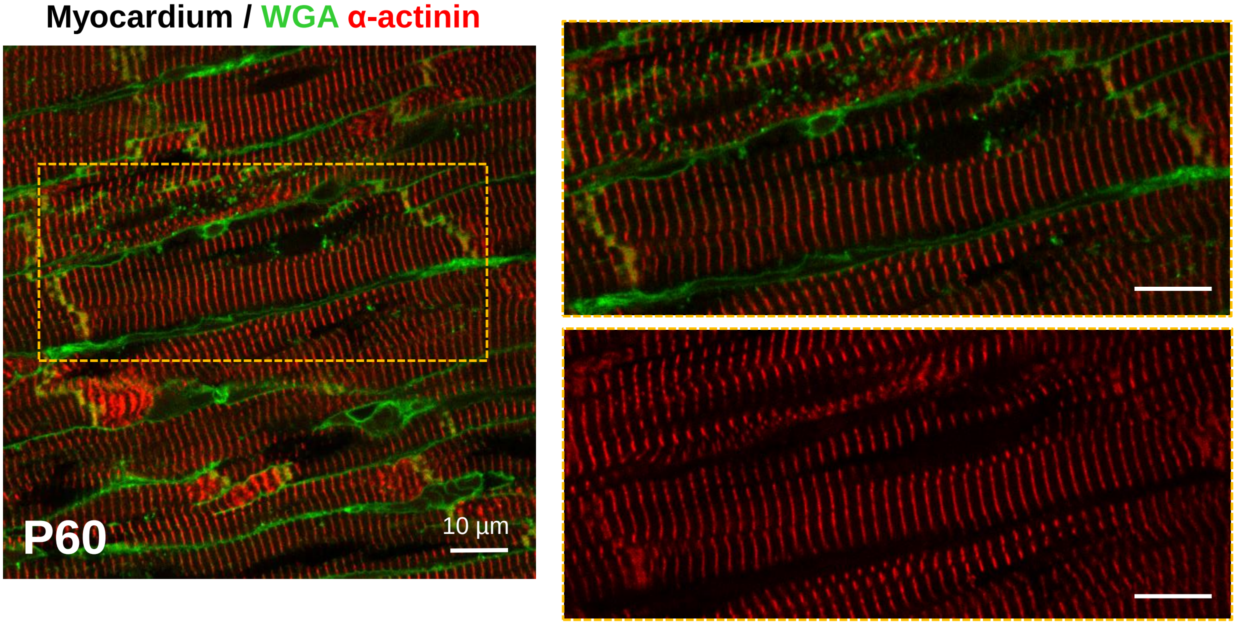
Myocardium / WGA α-actinin
P60
10 µm

*Figure V - Karsenty et al.*

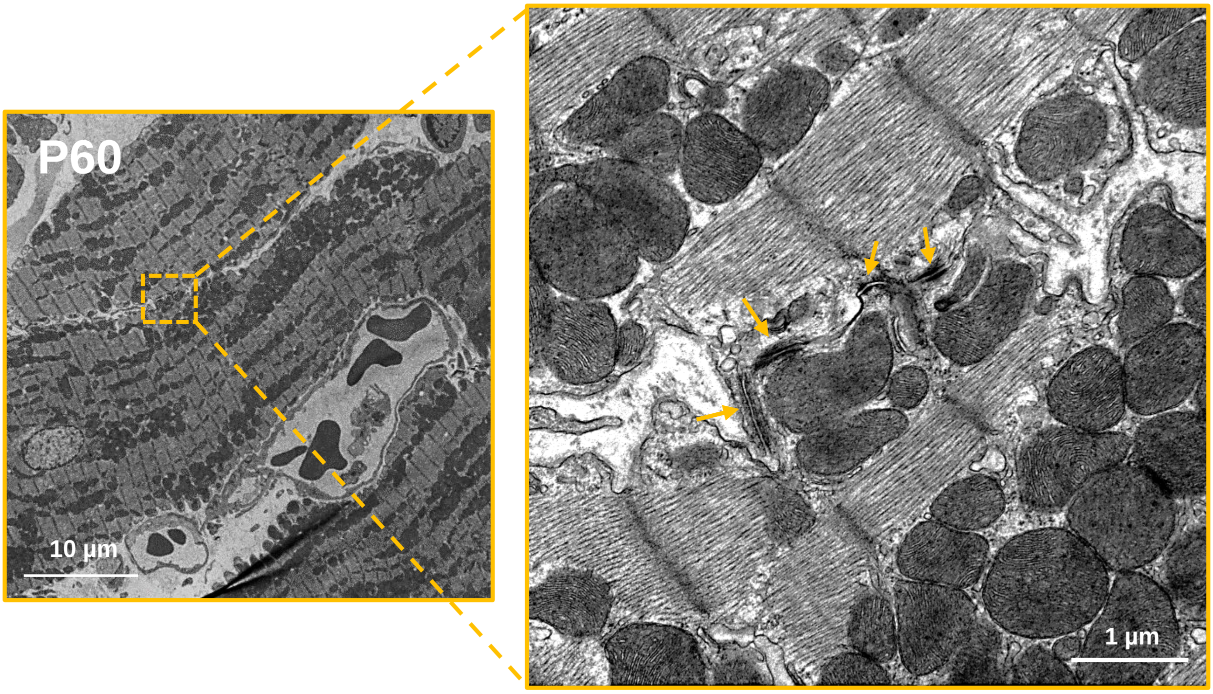

Figure VI- Karsenty et al.

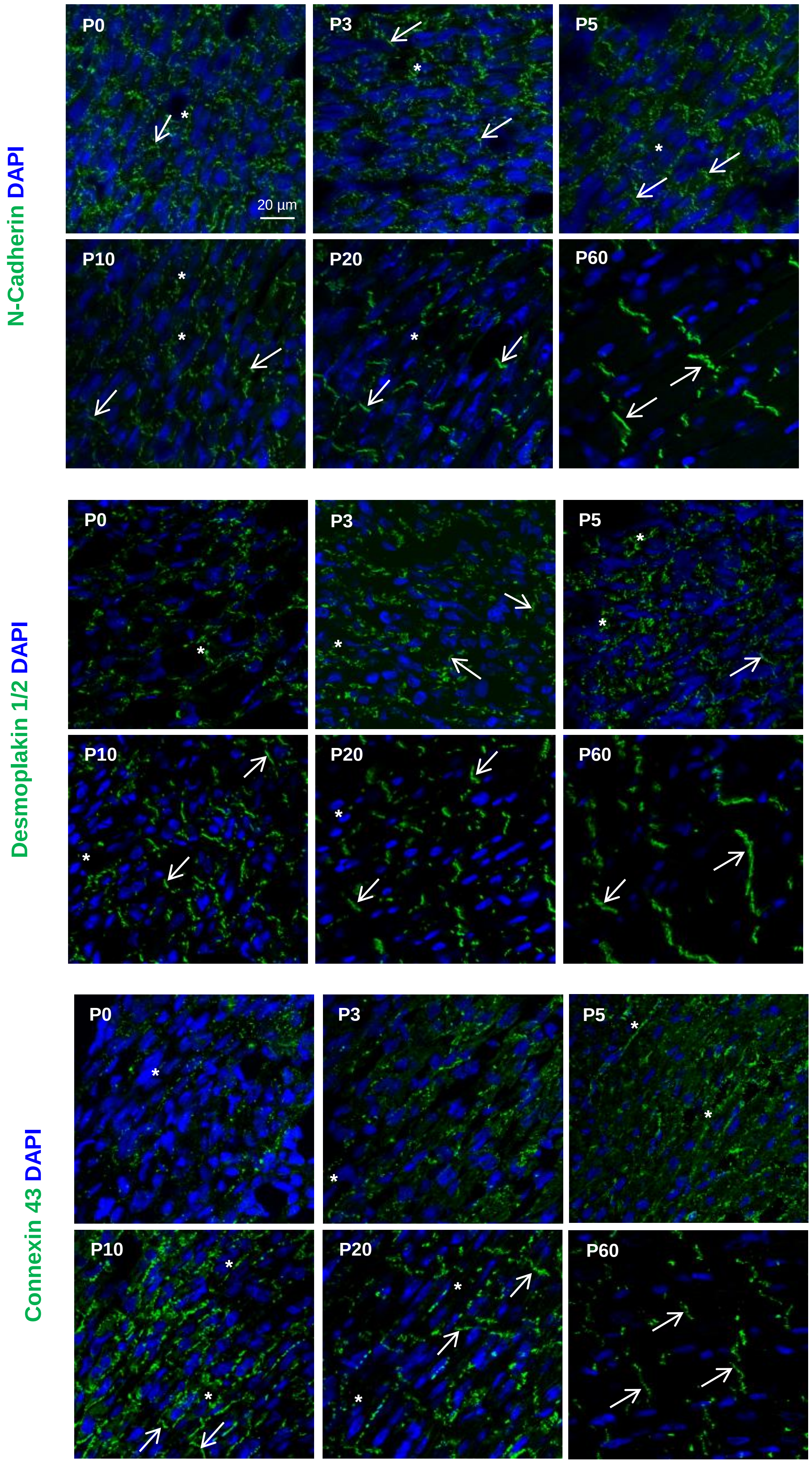

### Figure VII - Karsenty et al.

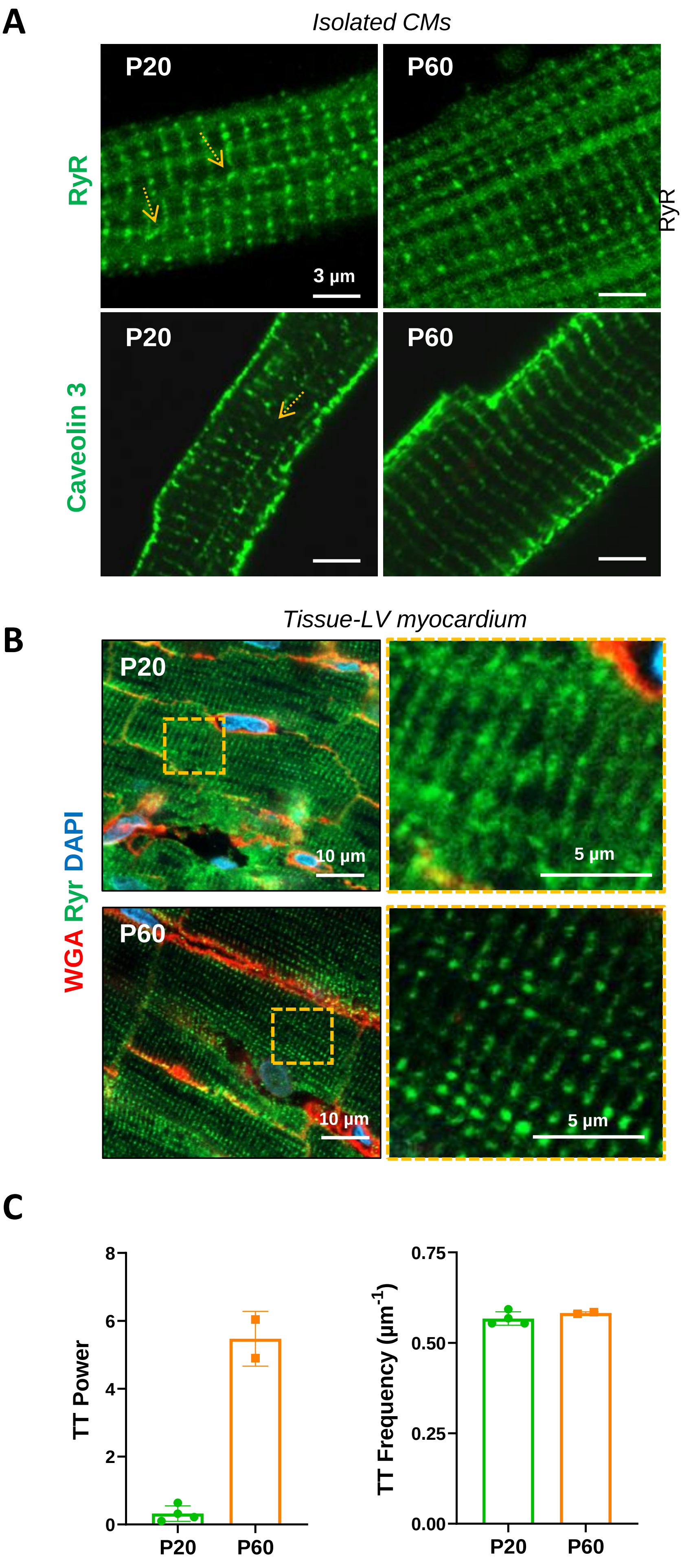

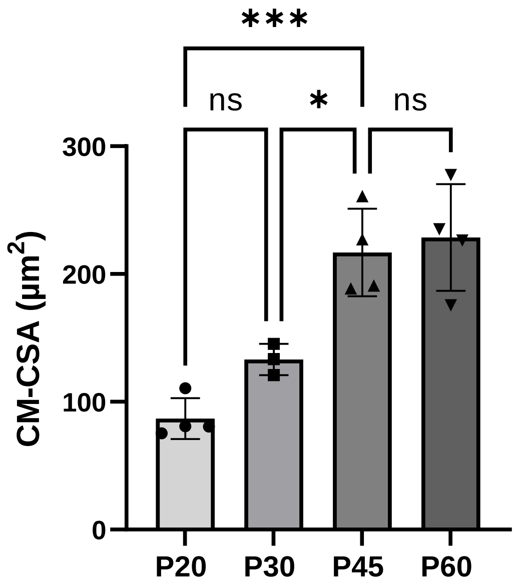
***
ns
*
ns
300
200
100
0
CM-CSA (µm2)
P20
P30
P45
P60

### Figure IX - Karsenty et al.

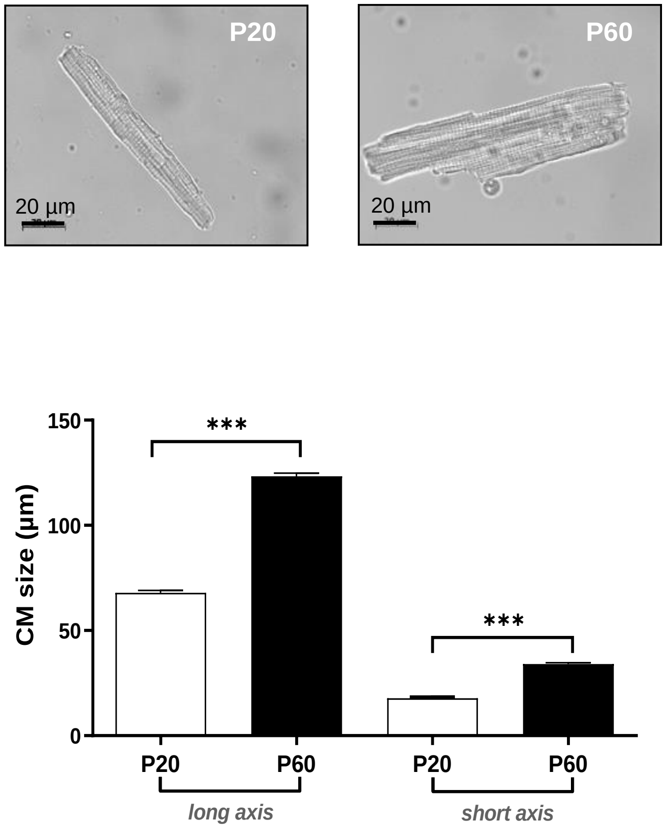

*Figure X- Karsenty et al.*

**A** Mice

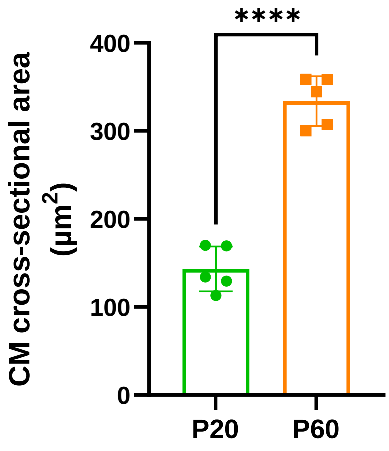


**B** Mice

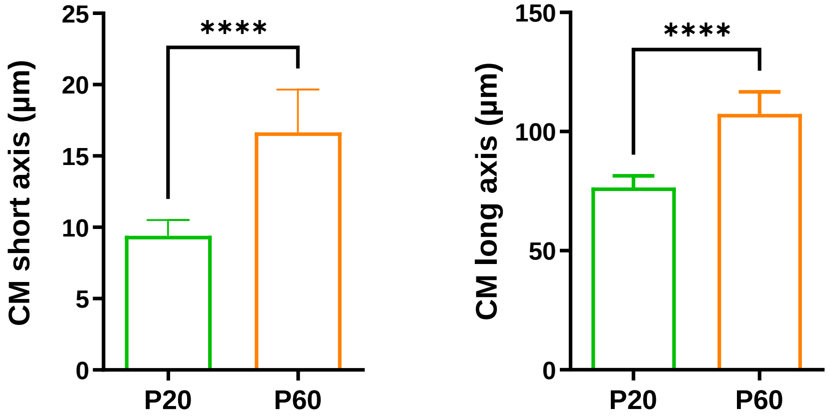

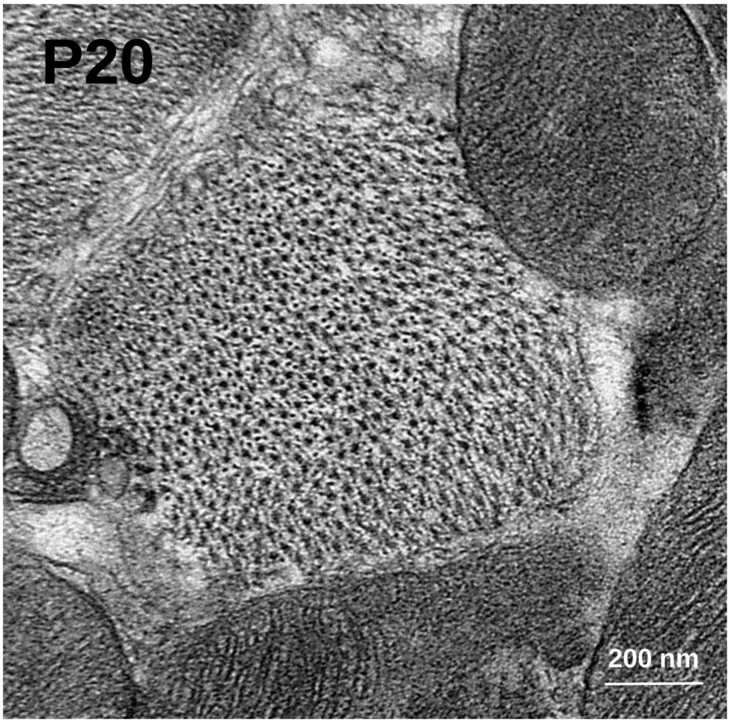
P20
200 nm

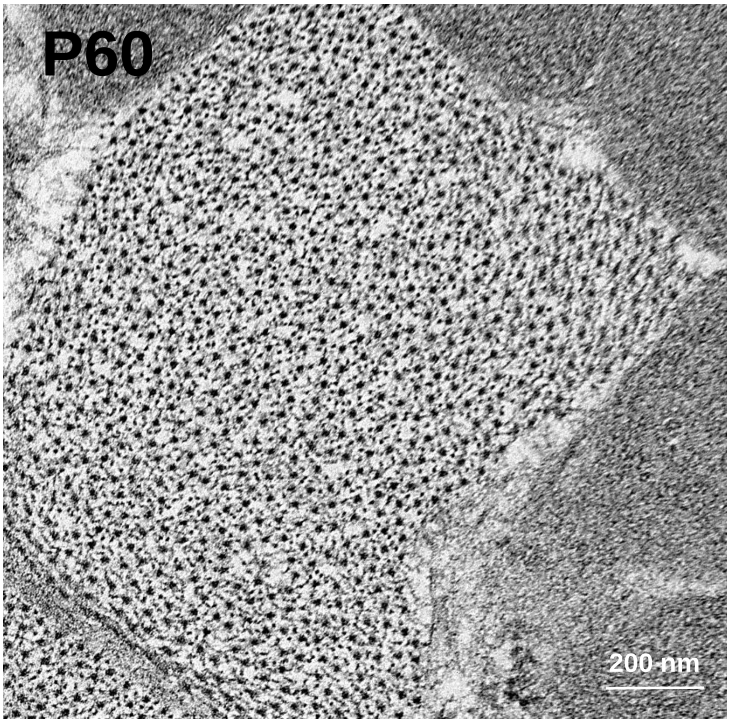
P60
200 nm

### *Figure XII - Karsenty et al.*

#### Up-regulated WT P60 vs P20 (482genes)

*GO categories*

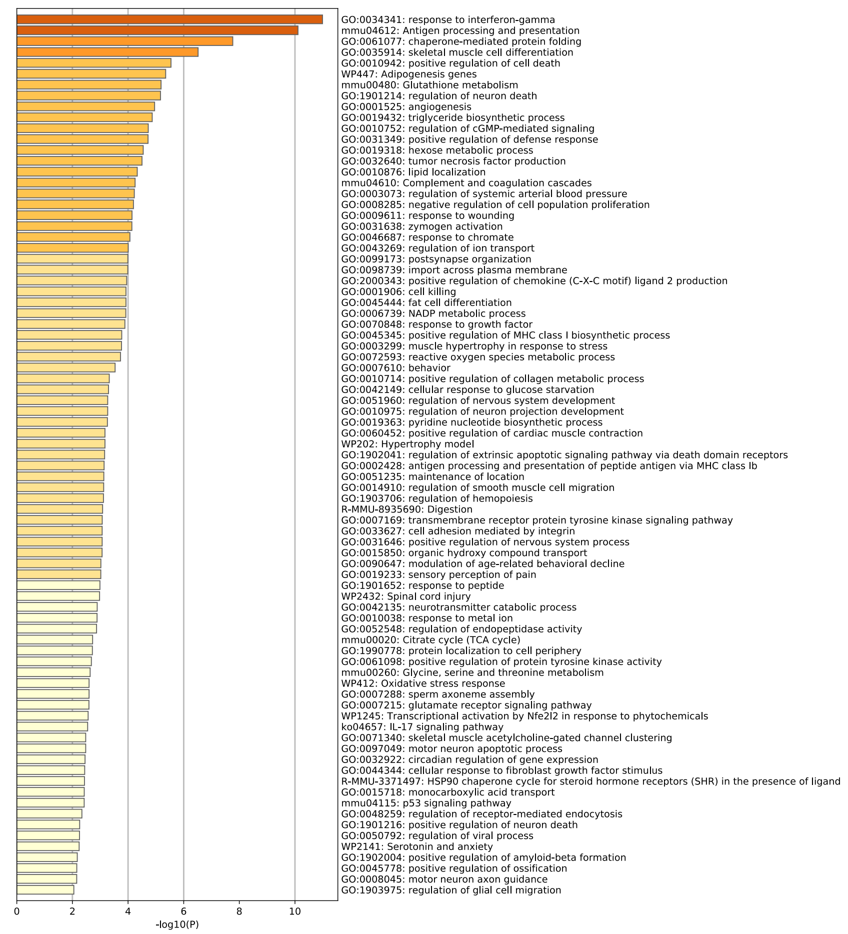


#### Down-regulated WT P60 vs P20 (464 genes)

*GO categories*

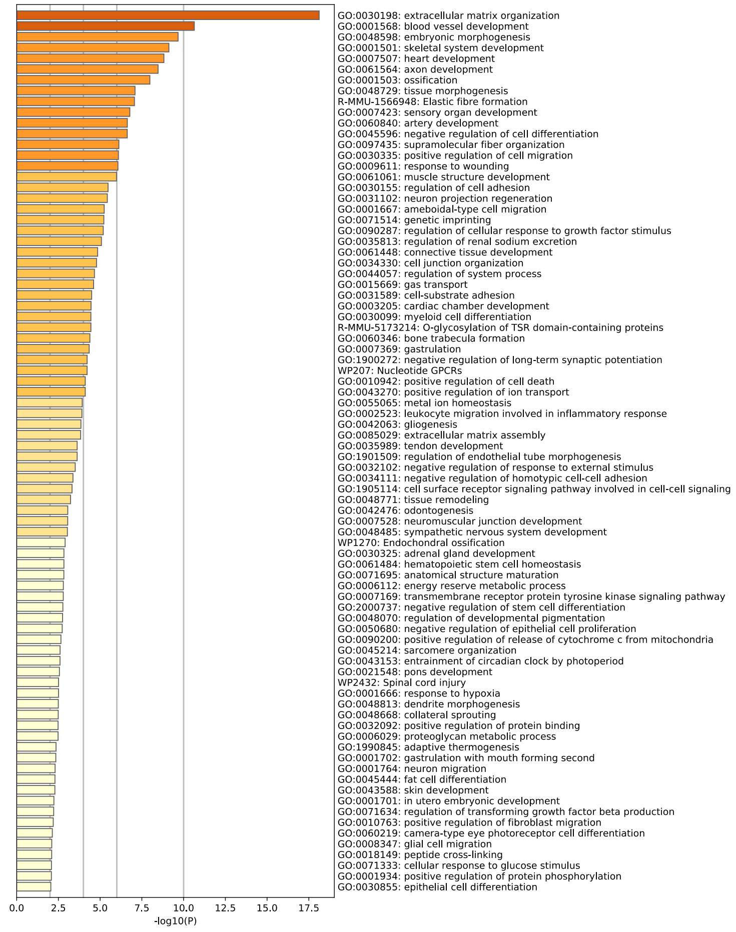

#### *Figure XIII - Karsenty et al.*

**A**

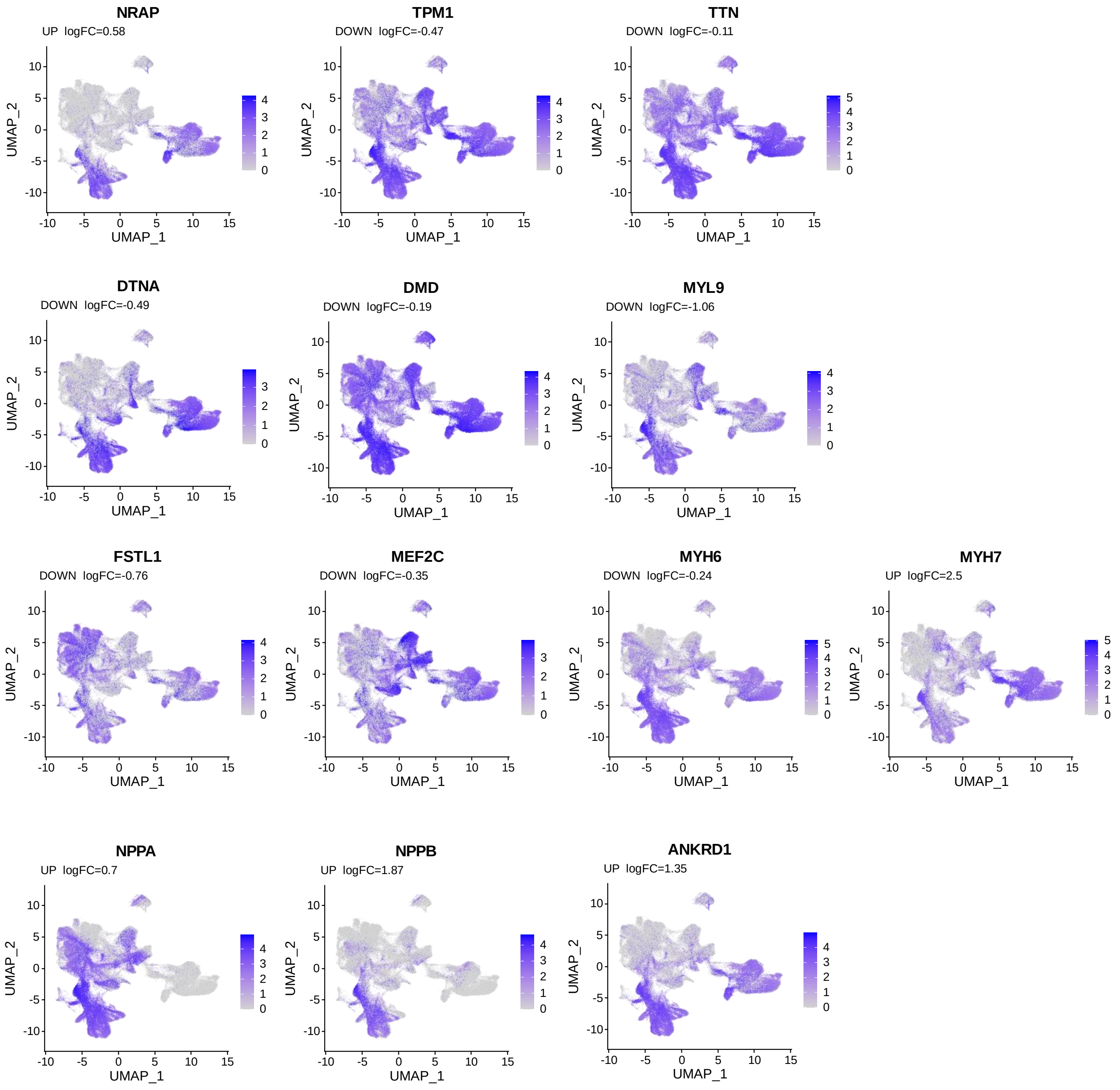


**B**

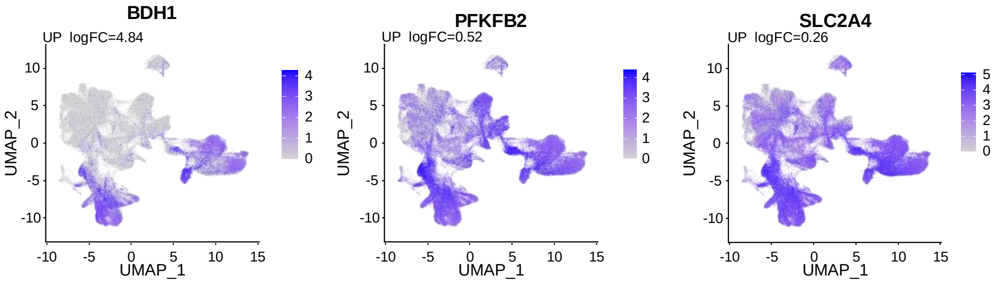

### *Figure XIV- Karsenty et al.*

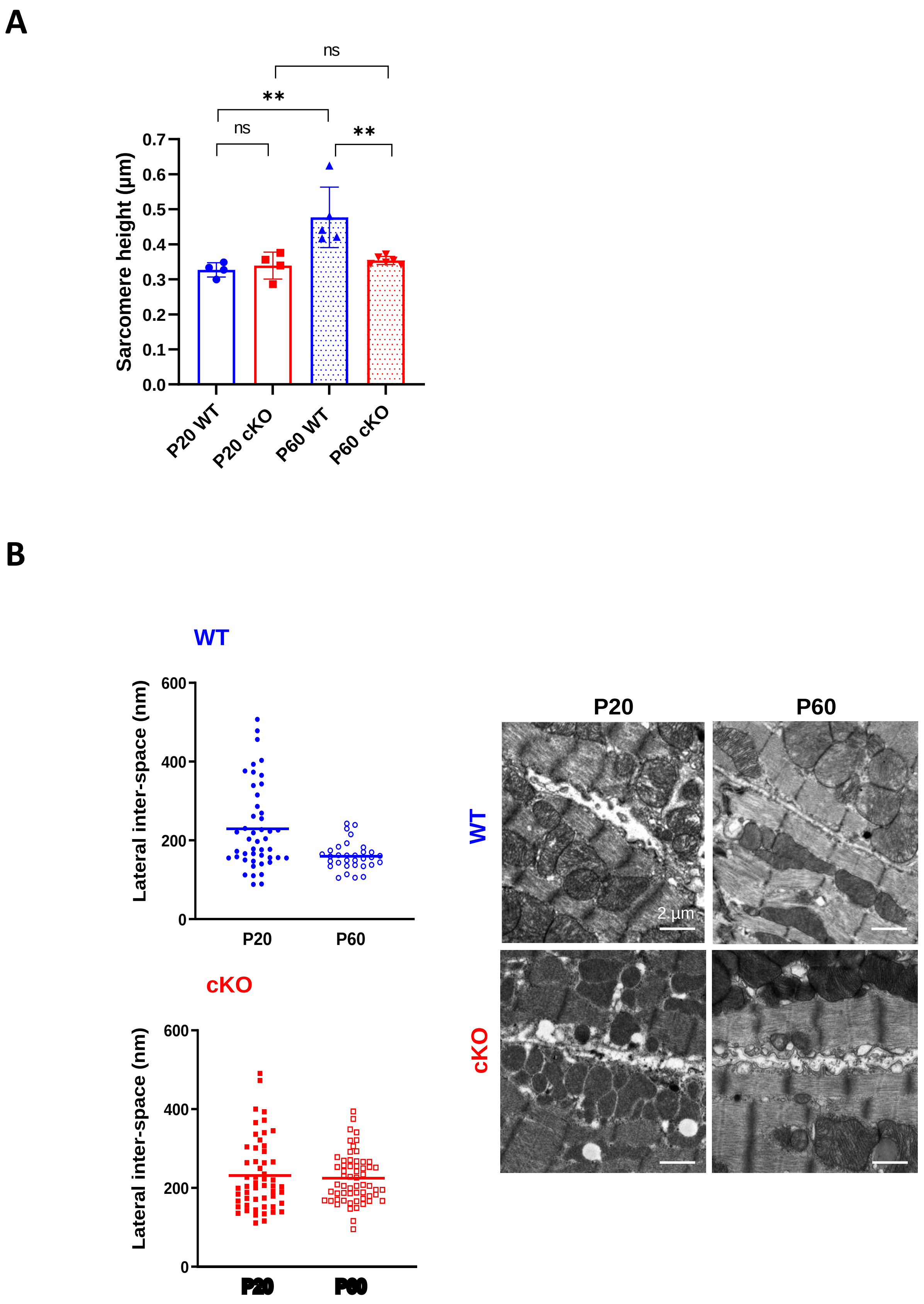
